# Coordinated host-pathogen transcriptional dynamics revealed using sorted subpopulations and single, *Candida albicans* infected macrophages

**DOI:** 10.1101/350322

**Authors:** José F. Muñoz, Toni Delorey, Christopher B. Ford, Bi Yu Li, Dawn A. Thompson, Reeta P. Rao, Christina A. Cuomo

**Affiliations:** Broad Institute of MIT and Harvard, Cambridge, MA USA; Worcester Polytechnic Institute, Worcester, MA USA

## Abstract

The outcome of fungal infections depends on interactions with innate immune cells. Within a population of macrophages encountering *C. albicans*, there are distinct host-pathogen trajectories; however, little is known about the molecular heterogeneity that governs these fates. We developed an experimental system to separate interaction stages and single macrophage cells infected with *C. albicans* from uninfected cells and assessed transcriptional variability in the host and fungus. Macrophages displayed an initial up-regulation of pathways involved in phagocytosis and proinflammatory response after *C. albicans* exposure that declined during later time points. Phagocytosed *C. albicans* shifted expression programs to survive the nutrient poor phagosome and remodeled the cell wall. The transcriptomes of single infected macrophages and phagocytosed *C. albicans* revealed a tightly coordinated shift in gene expression co-stages and revealed expression bimodality and differential splicing that may drive infection outcome. This work establishes an approach for studying host-pathogen trajectories to resolve heterogeneity in dynamic populations.

## Introduction

Interactions between microbial pathogens and the host innate immune system are critical to determining the course of infection. Phagocytic cells, including macrophages and dendritic cells, are key players in the recognition of and response to fungal infections^1^. *Candida albicans*, the most common fungal pathogen, can cause life threatening systemic infections in immunocompromised individuals; however, in healthy individuals, *C. albicans* can be found as a commensal resident of the skin, gastrointestinal system, and urogenital tract^2^. In addition, *C. albicans* can withstand harsh host environments, including the macrophage phagosome, by regulating metabolic and cell morphology pathways^3,4^. While macrophages directly control fungal proliferation and coordinate the response of other immune cells, the outcomes of these interactions are heterogeneous; some *C. albicans* cells are effectively killed by macrophage engulfment whereas others evade or survive macrophage interactions and persist in the host^5^.

Previous studies of *C. albicans* and immune cell interactions in bulk populations have identified key pathways by characterization of either the fungal or host transcriptional response during these interactions^3,6,7^. More recently, dual transcriptional profiling of host-fungal pathogen interactions has also examined populations of cells^8–11^. Bulk approaches measure the average transcriptional signal of millions of cells, obscuring differences between infection fates. Even in a clonal population of phagocytes, many immune cells do not engulf any fungal cells, while others can phagocytose up to ten fungal cells^12^. Single-cell RNA sequencing (scRNA-Seq) has highlighted the substantial variation in gene expression between cells within stimulated or infected immune cell populations^13–15^. For example, scRNA-Seq revealed that a subset of macrophages exposed to bacterial stimuli displayed a strong interferon-response, which was associated with cell surface variation between different bacteria^14^. A recent study measured host and pathogen gene expression in single host cells infected with the bacteria *Salmonella typhimurium;* however, the low number of pathogen transcripts detected per cell was only sufficient for analysis of sets of co-regulated genes^16^. To date, parallel transcriptional profiling of single host cells and fungal pathogens has not been reported.

To overcome these challenges, we developed an experimental system to isolate subpopulations of distinct infection outcomes and examined host and pathogen gene expression in sorted subpopulations and in single, infected macrophages. We focused on four distinct infection outcomes: (i) infected macrophages with live *C. albicans*, (ii) infected macrophages with dead, phagocytosed *C. albicans*, (iii) macrophages exposed to *C. albicans* that remained uninfected and (iv) *C. albicans* exposed to macrophages that remained un-engulfed. In addition to carrying out dual RNA-Seq on these subpopulations, we isolated single macrophages infected with *C. albicans* and adapted methods to measure gene expression of the host and pathogen to further resolve heterogeneity. By comparing the transcriptional profiles of *C. albicans* and primary, murine macrophages at both the subpopulation and single infected cell levels, we characterized the tightly coupled time-dependent transcriptional responses of the host and pathogen across distinct infection fates. We established that both host and pathogen gene expression can be measured from single cells; this revealed that genes involved in host immune response and in fungal morphology and adaptation show expression bimodality or changes in splicing patterns, variation that is important to consider in monitoring the dynamics of host-fungal pathogen interactions.

## Results

### Characterization of heterogeneous subpopulations *ex vivo* during macrophage and *Candida albicans* interactions

To capture infection subpopulations and more finely examine host and pathogen interactions, we developed a system for fluorescent sorting of *Candida albicans* with macrophages. We utilized a reporter to measure fungal cell status (live or dead) and infection status (engulfed or unengulfed). This construct, which constitutively expresses Green Fluorescent Protein (GFP) and mCherry, was integrated into *C. albicans* at the *NEUT5* locus (**Methods**); when *C. albicans* cells lyse in the acidic macrophage phagosome, GFP loses fluorescence upon the change in pH^15^ whereas mCherry remains stable for up to 4 hours in this environment as visualized by microscopy. Primary, murine bone derived macrophages were stained with CellMask Deep Red plasma membrane stain. To study host-fungal pathogen infection stages at finer resolution, the *C. albicans* reporter strain was then co-incubated with primary bone-derived, stained macrophages and subpopulations were isolated using fluorescence-activated cell sorting (FACS) at time intervals (0,1, 2 and 4 hours; **Methods; Figure 1A**). These time points were selected to capture the early transcriptional changes of *C. albicans* in response to interactions with macrophages^3^. To examine gene expression, RNA of both host and fungal cells was extracted and adapted for Illumina sequencing using Smart-Seq2 (**Methods**). Four major infection subpopulations were isolated by FACS: (i) macrophages infected with live *C. albicans* (GFP+, mCherry+, Deep red+), (ii) macrophages infected that phagocytosed and killed *C. albicans* (GFP-, mCherry+, Deep red+), (iii) macrophages exposed to *C. albicans* (GFP-, mCherry-, Deep red+) and (iv) *C. albicans* exposed to macrophages (GFP+, mCherry+, Deep red-; **Figure 1A**). The number of uninfected macrophages exposed to *C. albicans* ranged from an average of 61% to 67%, while the number of uninternalized *C. albicans* ranged from 22% to 7% over the time course (**Figure 1B**). The number of infected macrophages varied from 11% to 29% over the time course; the largest increase in this population was observed between 0 and 1 hours post infection and then remained stable over 2 and 4 hours (**Figure 1B**). The number of macrophages infected with dead, phagocytosed *C. albicans* ranged between 0% to 3% over the time course (**Figure 1B**).

**Figure 1.**
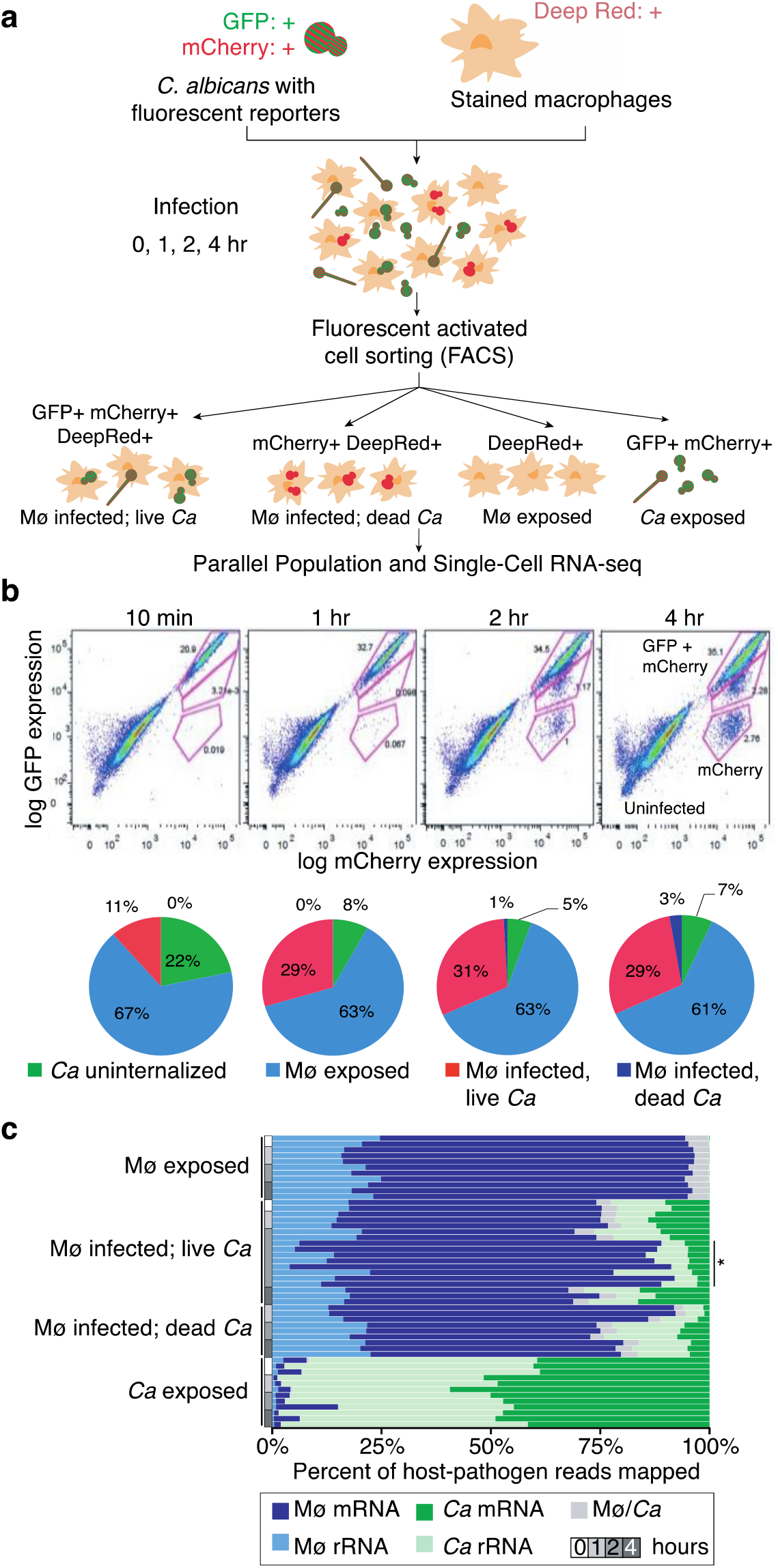
Heterogeneous populations of macrophage-Candida encounters are captured by cell sorting sequencing and expression analysis. **(a)** Schematic representation of the experimental model, using primary, bone marrow-derived macrophages (BMDMs) incubated with *Candida albicans* reporter strain CAI4-F2-mCherry-GFP, sampling at time intervals, and sorting to separate subpopulations of interacting cells. **(b)** Bone marrow derived mouse macrophages (Mø) were incubated with the *Candida albicans* (*Ca*) reporter strain (SC5314 Neut5L-NAT1-mCherry-GFP), then sorted at 10, 60, 120 and 240 minutes using fluorescence-activated cell sorting on the BDSORP FACSAria. Pie charts depicting the percent of cells sorted for each infection subpopulation. **(c)** Percent of host (Mø: macrophages) and pathogen (Ca: *C. albicans*) RNA-Seq reads mapped to the composite reference transcriptome, including mouse (GRCm38/mm10; mRNA and rRNA transcripts) and *Candida albicans* (CAI4-F2; mRNA and rRNA transcripts). Mø/Ca (gray color label) is the proportion of multiple mapped reads, *i.e*. reads that map to both the mouse and *C. albicans* transcriptomes, which were excluded from further analysis.

The number of RNA-Seq reads and transcripts detected for both host and fungal pathogen subpopulations was sufficient to profile parallel transcriptional responses (**Supplementary Note 1**). Based on alignments to a composite reference of both mouse and *C. albicans* transcriptomes (**Methods**), the fraction of mapped reads for host and pathogen was highly correlated with the size of the transcriptomes and percent of sorted cells for each subpopulation (*e.g*. 87% host and 13% fungus for macrophages infected with live *C. albicans;* **Figures 1C, S1B; Table S1**). In subpopulations of macrophages infected with live fungus, an average of 10,333 host and 4,567 *C. albicans* genes were detected (at least 1 fragment per replicate across all samples; **Figure S1A; Table S1**). We focused the differential expression analysis on subpopulations with high transcriptome coverage and highly correlated biological replicates (*e.g*. Pearson’s *r* > 0.85, and > 6,000 and 2,000 transcripts detected in macrophages and *Candida*, respectively; **Figure S1B**). The detection of a high number of transcripts across sorted subpopulations supports that we have established a robust system for defining host and fungal pathogen gene expression profiles during phagocytosis.

Next, we examined the major expression profiles in both the host and fungus. In *C. albicans* and macrophages, respectively, we identified 588 and 577 differentially expressed genes (DEGs; fold change (FC) > 4; false discovery rate (FDR) < 0.001) among all pairwise comparisons (**Methods**; **Figure 2**; **Data set 1** and **2**). To determine the major patterns of infection-fate specific or interaction-time specific, we used DEGs to perform Principal component analysis (PCA), then PC scores were clustered by *k-means*(**Methods**). PC analysis revealed that the transcriptional response in infected and exposed macrophages primarily varied over time and were highly similar at each time point between these subpopulations (**Figure 2C**). While exposed and phagocytosed *C. albicans* subpopulations also varied over time, these two populations appeared more distinct as described below (**Figure 2A**). These findings highlight how cell sorting can be used to separate different infection fates and distinguish gene expression signatures within populations of host and pathogen cells.

**Figure 2.**
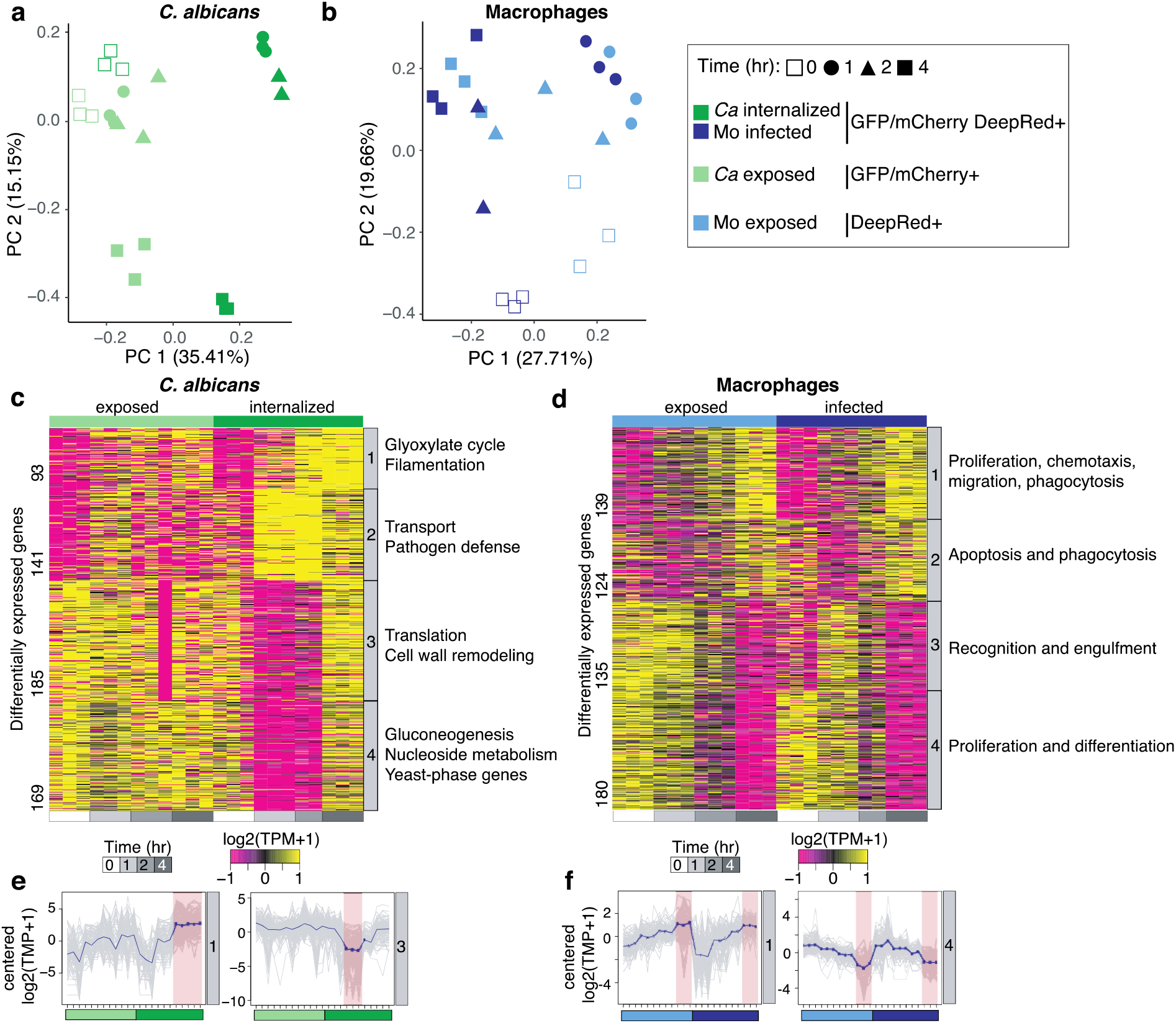
Subpopulations of infection outcome reveal dynamic parallel host-pathogen orchestration of transcriptional response. Principal component analysis (PCA) using the transcript abundance for the pathogen (**a**; *C. albicans*) and the host (c; macrophages) using significantly differentially expressed genes (DEGs; FC >4, FDR < 0.001) in live *C*. albicans-phagocytized macrophages (GFP+ mCherry+ Deep Red+) compared to all other conditions. Contributions of each infection outcome replicate (points) to the first two principal components (PC1 and PC2) are depicted. The projection score (red: high; blue: low) for each gene (row) onto PC1 and PC2 revealed four clusters in both host and pathogen. For each projection score cluster, immune response genes (macrophages) and functional biological categories (*C. albicans;* deducted from GO term enrichment analysis) are shown. (**b**) Transcriptional response in subpopulations of *Candida albicans*. Heatmap depicts significantly differentially expressed genes for replicates of each *C. albicans* sorted populations (exposed and phagocytosed) at 0, 1, 2 and 4 hours post-infection, grouped by *k-means* (similar expression patterns) in five clusters. Each cluster includes synthesized functional biological relationships using Gene Ontology (GO) terms (corrected P < 0.05). **(d)** Transcriptional response in subpopulations of macrophages. Heatmap depicts significantly differentially expressed genes (DEGs; FC >4, FDR < 0.001) for replicates of each macrophage-sorted populations (exposed and infected) at 0, 1, 2 and 4 hours post-infection, clustered by *k-means* (similar expression patterns). Each cluster includes synthesized functional biological relationships using IPA terms (-log(p-value) > 1.3; z-score > 2). Expression patterns for **(f)** clusters 1 and 4 in *C. albicans* and **(g)** cluster 1 and 3 in macrophages. Each gene is plotted (gray) in addition to the mean expression profile for that cluster (blue).

### Subpopulations of phagocytosed *C. albicans* adapt to macrophages by switching metabolic pathways and regulating cell morphology

We next examined how *C. albicans* gene expression varied across exposed and phagocytosed cells over time. Using *k-means* clustering of 588 DEGs, we identified 4 clusters of genes with similar expression patterns; the major patterns of expression across time were either induced rapidly upon phagocytosis (clusters 1 and 2) or repressed at 1 and 2 hours upon phagocytosis (clusters 3 and 4; **Figures 2B, S2A**). Comparing the phagocytosed and un-engulfed *C. albicans* subpopulations at each time point, the highest number of DEGs was found at 1 hour (n = 165), highlighting a rapid and specific transcriptional response upon phagocytosis, which was maintained throughout the 4-hour infection time course (**Table S2; Figure 2B**).

Genes highly induced in phagocytosed *C. albicans* at 2 and 4 hours (cluster 1, *n* = 93) are involved in adaptation to the macrophage environment including changes in metabolic pathways (**Figure 2B**). These genes are involved in glucose and carbohydrate transport, carboxylic acid and organic acid metabolism, and fatty acid catabolic processes (enriched GO terms corrected-P < 0.05, hypergeometric distribution with Bonferroni correction; **Table S3**). Prior microarray analysis of bulk populations of *C. albicans* exposed to macrophages reported similar changes^3^; by sorting infection fates, our work suggests that this response is specific to engulfed *C. albicans*, allowing the pathogen to utilize the limited spectrum of nutrients available in the phagosome^3^. We also found that genes involved in glyoxylate metabolism, the beta-oxidation cycle, and transmembrane transport were significantly induced in phagocytosed *C. albicans* relative to exposed cells. Using sorted populations revealed that multiple classes of transporters were highly up-regulated in both engulfed and exposed *C. albicans* subpopulations (including oligopeptide transporters, several high affinity glucose transporters, and amino acid permeases), suggesting these changes are not in response to phagocytosis (**Figure 2B; Tables S2** and **S3**). Genes most strongly induced during at 4 hours upon phagocytosis (cluster 1) are involved in pathogenesis and associated with the formation of hyphae, including four of the eight genes involved in the core filamentation response^17^ (*ALS3, ECE1, HWP1, orf19.2457;* **Table S2**). While media containing serum can also induce *C. albicans* filamentation, we found that filamentation genes were more highly induced in the phagocytosed *C. albicans* subpopulation (**Table S2; Figure S2B**). Genes induced most strongly at 1 and 2 hours upon phagocytosis (cluster 2) are involved in early fatty acid oxidation response and transmembrane transport (Opt and Hgt classes; Tables S2 and S3). These transporters differ from those in cluster 1 in that they show peak expression at 1 hour in phagocytosed cells and decrease in expression by 4 hours (**Figure 2B**). Clusters 1 and 2 contained 6 confirmed or putative transcription factors (*SUT1, STP4, TEA1, ADR1, ZCF38, TRY4*), all of which encode zinc finger containing proteins; zinc cluster transcription factors have been implicated in *C. albicans* virulence^18,19^.

Two sets of genes were specifically down-regulated in live, phagocytosed *C. albicans* at 1 and 2 hours. Expression levels of these genes largely did not change in exposed *C. albicans* over the time course (clusters 3 and 4; **Figures 2B, 2E**). Notably, these repressed genes recovered their expression levels by 4 hours in phagocytosed *C. albicans*. Cluster 3 included genes related to the translation machinery and peptide biosynthesis, including ribosomal proteins, chaperones and transcription factors that regulate translation (enriched GO terms, corrected-P < 0.05, hypergeometric distribution with Bonferroni correction; **Table S3; Figure S2B**). Repression of the translation machinery was previously noted in *C. albicans* during macrophage interaction^3^. Our results demonstrate that down-regulation of ribosomal proteins, chaperones and translation-regulator transcription factors are specific to phagocytosed *C. albicans* and that expression of these genes recovered at later time points (**Figures 2E, S2B**). Cluster 3 also encompassed genes involved in morphological and cell surface remodeling, including an essential negative regulator of filamentation, *SSN6* (**Figures 2B, S2B; Table S2**). Repressed genes in Cluster 4 are largely involved in nucleoside metabolic processes, gluconeogenesis and host adaptation (enriched GO terms, corrected-P < 0.05, hypergeometric distribution with Bonferroni correction; **Figure 2B; Table S3**). A subset of these repressed genes are yeast-phase specific and included those involved in ergosterol biosynthesis, cell growth and cell wall synthesis (**Figures 2B, S2**). These results highlight that a large part of the observed transcriptional repression is specific to phagocytosed *C. albicans*, in contrast to the common sets of up-regulated genes shared by exposed and phagocytosed cells.

### Subpopulations of macrophages showed major pathogen recognition and pro-inflammatory response to *C. albicans* and shift profiles at late time course

In parallel with the analysis of *C. albicans* gene expression, we also examined the transcriptional response of macrophages. Across all samples, we identified 577 DEGs (FC > 4; FDR < 0.001; Data set 2), which grouped into four clusters with similar expression patterns (**Figures 2C, 2D**). For both exposed and infected macrophages, a major difference was found between 1 and 4 hours along PC1, which highlights genes that were highly induced or repressed at 4 hours in these subpopulations (clusters 1 and 4, respectively; **Figure 2C, D**). These 4 clusters together were significantly enriched for genes involved in activation of phagocytosis, migration of phagocytes, and triggering the innate immune response; this includes the induction of pathways such as IL-6, IL-8 and NF-κB signaling, Fcy Receptor-mediated phagocytosis, production of nitric oxide and reactive oxygen species (ROS), Pattern Recognition Receptors (PRR), RhoA, ILK, and leukocyte extravasation signaling (P-value < 0.05 right-tailed Fisher’s Exact Test; **Figures 2D, S3**). Activation of some of these pathways is consistent with previous analysis of phagocyte transcriptional responses to *C. albicans* infection (**Figure S4**)^8–10,20^; sorting distinct infection subpopulations during early time points of infection established that host cells regulate subsets of genes upon *C. albicans* exposure and phagocytosis (**Figures 2D, S5**).

In exposed and infected macrophages, many of the genes induced at 1 hour maintained this expression level at 2 and 4 hours (cluster 1; **Figure 2D**). These genes are related to defense mechanisms such as pro-inflammatory cytokine production and fungal recognition via transmembrane receptors. Up-regulated genes related to pro-inflammatory cytokines included tumor necrosis factor (*Tnf*), interferon regulatory factor 1 (*Irf1*), and the chemokine receptor *Cx3cr1*, associated with an innate mechanism of fungal control in a model of systemic candidiasis^21^ and colitis^22^ (**Figures 2D, S5; Table S4**). A second set of genes was initially repressed at 1 hour and then up-regulated in both exposed and infected macrophages at 2 and 4 hours (cluster 2; **Figure 2D; Table S4**). This set of genes is associated with pathogen recognition, opsonization, and activation of the engulfment (P-value < 0.05 right-tailed Fisher’s Exact Test; **Table S5**), including the lectin-like receptor galectin 1 (*Lgals1*), transmembrane receptors (*Fcer1g*), chemokines (*Ccl3, Cxcl2*), extracellular complement protein (*C1qb*), and transcriptional regulators that play a role in inflammation and programmed cell death (*Fos, Irf8, Cebpb* and *Card9*) (**Tables S4 and S5**). Since the expression of these genes increased at 2 hours and maintained high expression in infected macrophages, they may also play an important role during phagocytosis or allow for uptake of additional *C. albicans* cells. The chemokines *Cxcl2, Ccl3* and *Cx3cr1* have also been previously shown to be induced during *C. albicans* interactions with other host cells, including neutrophils *in* vitro^10^, in a murine kidney model^8^, a murine vaginal model^20^, and in mouse models of hematogenously disseminated candidiasis and of vulvovaginal candidiasis in humans^9^, highlighting the role of these genes in host defense against *C. albicans* infection of different tissues (**Figure S4**).

We also examined subpopulations of macrophages that have phagocytosed and killed *C. albicans;* this data was analyzed separately, as the total number of cells sorted and therefore the transcriptome coverage were low (< 3,000 host transcripts detected) and had modestly correlated biological replicates (*e.g*. Pearson’s *r* < 0.56). We found a small set of highly induced pro-inflammatory cytokines, including *Ccl3, Cxcl2, Il1rn* and *Tnf*, and transcription regulators such as *Cebpb, Irf8* and *Nfkbia* (Figure S6). These genes were also induced in macrophages infected with live *C. albicans* (clusters 1 and 2; **Figure 2D**), indicating that maintaining expression of these genes may be important for pathogen clearance and host cell survival after phagocytosis.

Another major shift in macrophage gene expression occurred at 4 hours, with sets of genes involved in the immune response highly repressed at this later time point (clusters 3 and 4, respectively; **Figures 2D, 2F**). Repressed genes at 4 hours (cluster 3) are enriched in cytokines (*Il1a, Cxcr4*) and transmembrane receptors, including intracellular toll-like receptor (Tlr5), C-type lectin receptors (*Clec4a3, Clec10a, Olr1*), and interleukin receptors (Il1r1). This cluster was also enriched in categories associated with proliferation and immune cell differentiation (P-value <0.05 right-tailed Fisher’s Exact Test; **Table S5**). In addition, highly repressed host genes at 4 hours (cluster 4) were enriched for categories related to phagosome formation, phagocytosis signaling, and immune response signaling (P-value < 0.05 right-tailed Fisher’s Exact Test; Table S5). Notably, these repressed genes included several interleukin receptors (*Il21r, Il4r, Il17ra*) and transmembrane receptors (*Tlr9, Mrc1*) that are typically highly expressed during the immune response to fungal infections^8,10^. This suggests that during phagocytosis of *C. albicans* there is a strong shift in macrophages toward a weaker pro-inflammatory transcriptional response by 4 hours.

### Detection of host-pathogen transcriptional responses from single macrophages infected with *C. albicans*

Even in sorted populations, individual cells may not have uniform expression patterns, as, even in a clonal population, cells can follow different trajectories over time. To address this, we next examined the level of single cell transcription variability during these infection time points. We collected sorted, single macrophages infected with live or with dead, phagocytosed *C. albicans* at 2 and 4 hours and adapted the RNA of both the host and pathogen for Illumina sequencing using Smart-Seq2 (**Figure 1A; Methods**). With this approach, each infected macrophage and the corresponding phagocytosed *C. albicans* received the same sample barcode, allowing us to pair transcriptional information for host and pathogen at the single, infected cell level (**Figure 3A**). While we successfully isolated single infected macrophages via FACS, we cannot control for the number of *C. albicans* cells inside of each macrophage with this approach, since macrophages can phagocytose variable numbers of *C. albicans* cells^12^. We obtained 4.03 million paired-end reads per infected cell on average; a total of 449 single, infected macrophages had more than 1 million paired-end reads (**Table S1**). For macrophages with live *C. albicans*, we found an average 75% of reads mapped to host transcripts and 11% of reads mapped to *C. albicans* transcripts (**Figure 3B**). Although parallel sequencing of host and pathogen decreases the sensitivity to detect both transcriptomes from a single library, of the 224 single macrophages infected with live *C. albicans* with more than 0.5 million reads, 202 (90.2%) had at least 2,000 host-transcripts detected (> 1 Transcripts Per Million, TPM; 3,904 on average), and 162 (72.3%) had at least 600 *C. albicans* transcripts detected (> 1 TPM; 1,435 on average; **Figures 3B, S7**). The fact that we detected fewer fungal transcripts relative to the host was expected, as the fungal transcriptome is approximately four times smaller than the host transcriptome. Relative to the number of transcripts detected in the RNA-Seq of subpopulations of macrophages infected with live *C. albicans*, in single-infected-cells we detected 38% and 31% (on average) of the transcripts for host and *C. albicans* respectively, indicating that we obtained adequate sequencing coverage for both species. Additionally, we found that pooling single infected cell expression measurements could recapitulate the corresponding subpopulation expression levels. We found that the extensive cell-to-cell variation between single infected macrophages (average Pearson’s from *r* 0.18 to 0.88; **Figure 3C**) was reduced when we aggregated the expression of 32 single-cells (**Figure S8**). These results are consistent with previous single cell studies of immune cells^13,14,23^ and indicate that we can accurately detect gene expression in single infected macrophages and phagocytosed *C. albicans*.

**Figure 3.**
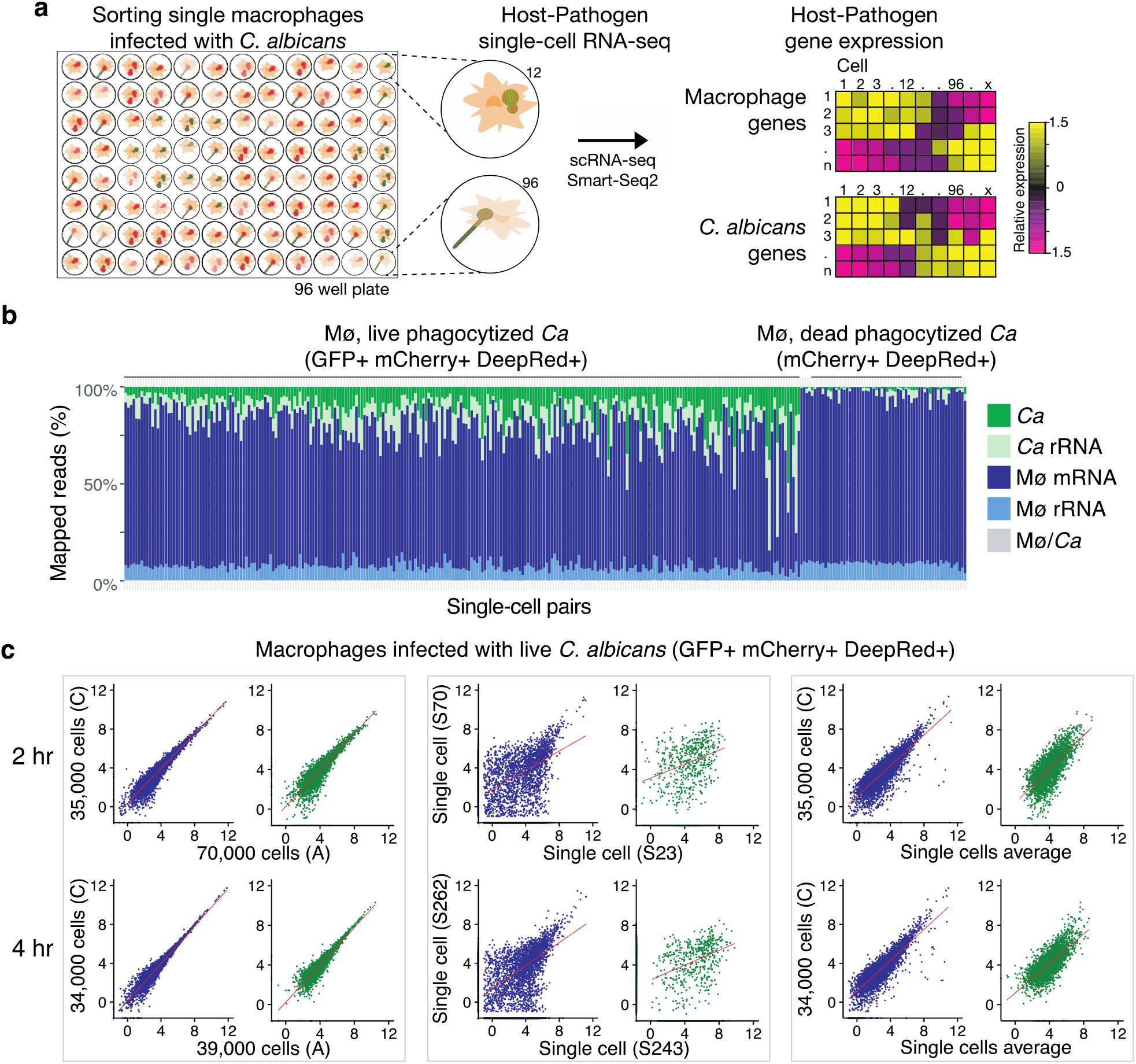
Processing and evaluation of parallel host-pathogen single-cell RNA-Seq. **(a)** Plot of the percent of mapped reads to the composite reference transcriptome (mouse messenger and ribosomal RNA +*Candida albicans* messenger and ribosomal RNA collections) of 314 single macrophages with live (mCherry + GFP+) or with dead, phagocytosed (mCherry + GFP-) phagocytosed *C. albicans* cells. **(b)** Plots of the gene expression correlation between (left) two replicates of the sorted subpopulation of macrophages (blue) with phagocytosed live *C. albicans* (green) at 2 and 4 hours post-infection; (middle) between two single macrophages and two phagocytosed *C. albicans*, and (right) between the single cells expression average and one replicate of the population.

### Dynamic host-pathogen co-stages defined by analysis of single macrophages infected with live *C. albicans*

To finely map the basis of heterogeneous responses during infection, we clustered cells by differential expressed genes and identified host-pathogen co-stages of infection in groups of infected macrophages and phagocytosed *C. albicans* pairs that showed similar gene expression profiles (**Figures 3A, 4A**). Briefly, we used genes exhibiting high variability across the infected macrophages and live, phagocytosed *C. albicans* at 2 and 4 hours. We then reduced the dimensionality of the expression with principal components analysis (PCA), and clustered cells with the t-distributed stochastic neighbor embedding approach (t-SNE)^24^ as implemented in Seurat^25^ (**Methods**). The transcriptional response among single infected macrophages exhibited two time-dependent stages associated with expression shifts from 2 to 4 hours (stage 1^M^ and stage 2^M^; **Figure 4A; Figure S9A**). While cells were largely separated into these two stages by time, a small subset of macrophages were assigned to the alternate cell stage by unsupervised clustering and appeared to be either early or delayed in the initiation of the transcriptional shift in genes involved in the immune response. Differentially expressed genes in stage 1^M^ (n = 88; likelihood-ratio test (LRT)^26^, *P* < 0.001) are related to the pro-inflammatory response, and their expression significantly decreases in stage 2^M^ at 4 hours (DEGs *n* = 70; LRT^26^, *P* < 0.001; Figure 4A, top; **Table S6**). This set comprises pro-inflammatory repertoire, such as cytokines (*Tnf, Ilf3, Ccl7*), transmembrane markers (*Cd83, Cd274*), interleukin receptors (*Il21r, Il4ra, Il6ra, Il17ra*) and the high affinity receptor for the Fc region (*Fcgr1*), and the transcriptional regulators *Cebpb* and *Cebpa* (**Figures 4A, 4B**). Many genes variably expressed in these single, infected macrophages were also found to be up-regulated in subpopulations of infected and exposed macrophages (*e.g. Tnf, Orl1*, **Figures S5, S10**); however, significant differences of these genes were not detected in the RNA-Seq of subpopulations between 2 and 4 hours, which highlights cell-to-cell variability within each time point that can only be observed in single infected macrophages.

As each single, infected macrophage received a unique sample barcode and host and fungal transcription were measured simultaneously (**Figure 3A**), the expression from each macrophage (**Figure 4A, top**) was matched with that of the live, phagocytosed fungus (**Figure 4A, bottom**). Notably, independent analysis of the parallel fungal transcriptional response identified two pathogen stages that were also primarily distinguished by time. In *C. albicans*, genes significantly up-regulated in stage 1^C^ (*n* = 80; LRT^26^, *P* < 0.001; **Table S7**) were enriched in organic acid metabolism (*P* = 6.63e-11; enriched GO term, corrected-P < 0.05, hypergeometric distribution with Bonferroni correction; **Table S8**), including transporters (*HGT13*) and glyoxylate cycle genes, specifically those from beta-oxidation metabolism (*ECI1, FOX3, FOX2, PXP2*; **Figures 4A, 4B**). Most macrophages infected by *C. albicans* in stage 1^M^ induced a strong pro-inflammatory response (co-stage of infection 1; **Figure 4**). At 4 hours in stage 2^C^, expression of these transporters and glyocylate cycle genes was reduced instead expression of genes (*n* = 86; LRT^26^, *P* < 0.001; **Table S7**) enriched in carbon metabolism was increased of (*P* = 2.43e-05; enriched GO term, corrected-P < 0.05, hypergeometric distribution with Bonferroni correction; **Table S8**), including genes related to glycolysis and gluconeogenesis (*PGK1*), fatty acid biosynthesis (*FAS1, ACC1*), and genes associated with filamentation (*ECE1, HWP1, OLE1, RBT1*); at this stage, the majority of infected macrophages down-regulated expression of pro-inflammatory cytokines (co-stage of infection 2; **Figure 4A**).

**Figure 4.**
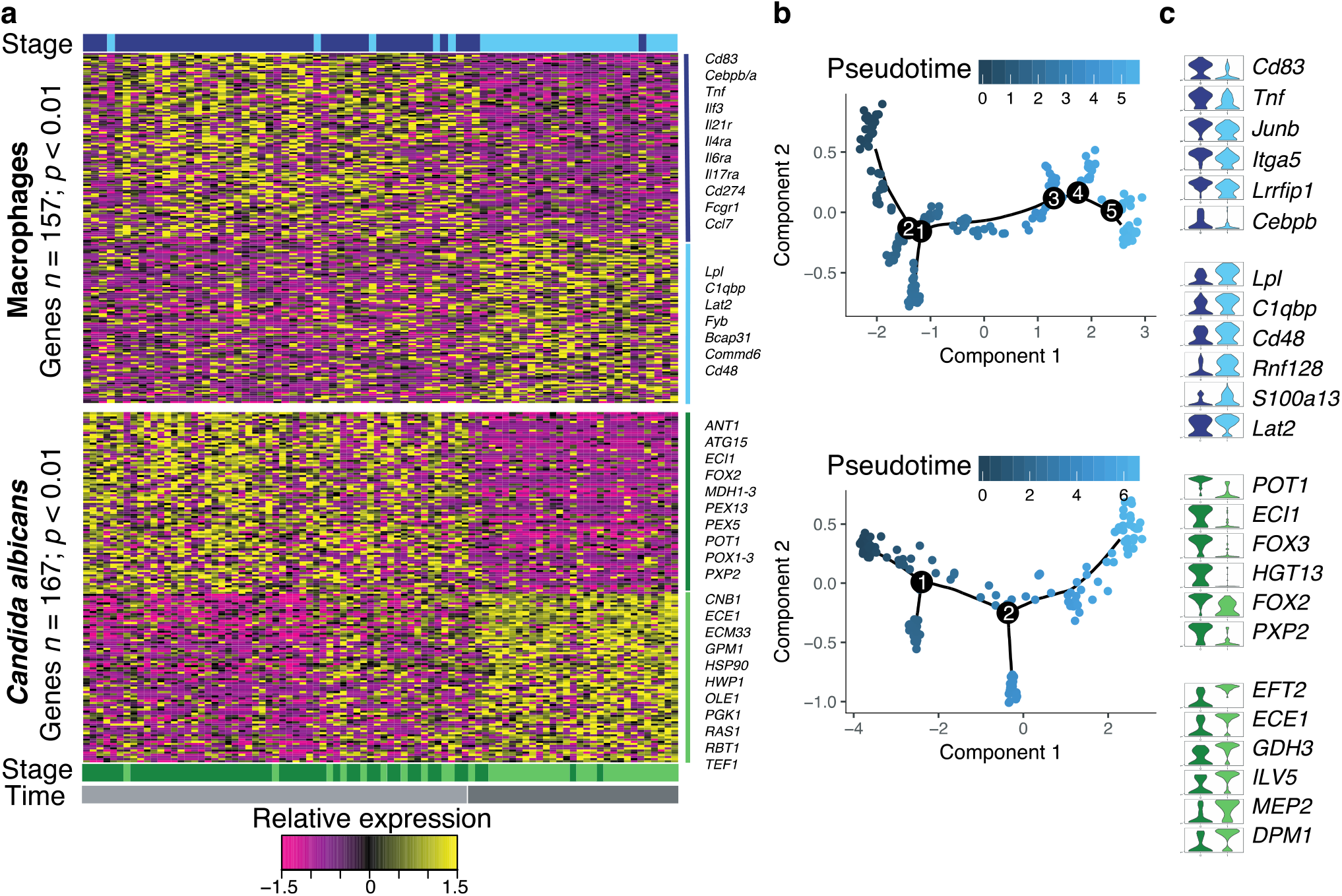
Parallel host-pathogen single-cell RNA-seq profiling reveals infection outcome shifts in pro-inflammatory response and pathogen immune evasion. **(a)** Heat maps report parallel scaled expression [log TPM (transcripts per million) +1 values] of differentially expressed genes for each costate of macrophages infected with live *C. albicans*. Each column represents transcriptional signal from a single, infected macrophage and the *C. albicans* inside of it. (Top: macrophage response; bottom: *C. albicans* response). Color scheme of gene expression is based on *z-score* distribution from −1.5 (purple) to 1.5 (yellow). Bottom and right margin color bars in each heat-map highlight co-state 1 (dark blue in macrophages, and dark green in *C. albicans*) and co-state 2 (blue in macrophages, and green in *C. albicans*), and time post-infection 2 hours (gray) and 4 hours (dark gray). **(b)** Violin plots at right illustrate expression distribution of a subset six differentially expressed immune response or immune evasion genes for each co-state in macrophages and *C. albicans*, respectively. **(c)** tSNE plot for macrophages (top) and *C. albicans* (bottom) colored by the pseudo-time.

To more finely trace how cells shift their expression program between infection co-stages we performed pseudo-time analysis using Monocle^27^ (**Methods**). We observed that the majority of infected macrophages and phagocytosed *C. albicans* pairs followed a linear expression trajectory between two major endpoints or clusters, recapitulating the two major infection co-stages; some cells exhibited alternative expression paths, suggesting the existence of minor cell trajectories (**Figures 4B, S11**). This ordering of cells revealed an early group of infected macrophages that expressed high levels of pro-inflammatory cytokines that decreased over time across intermediate and late groups (**Figures 4B, top; S11**). Similarly, in phagocytosed *C. albicans*, an initial group of cells showed low expression levels of genes related to filamentation which increased over time (**Figure 4B, bottom; S11**). In this analysis, the fraction of phagocytosed *C. albicans* shifting to high expression of filamentation genes across pseudo-time increased slightly faster than the fraction of macrophages exhibiting decreased levels of expression of pro-inflammatory cytokines (10% more cells in the late pseudo-time range; **Figure 4B**). This suggests that the expression heterogeneity in macrophages could be driven by the expression heterogeneity in *C. albicans* (**Figure 4B**). This was also supported by unsupervised clustering, where some phagocytosed *C. albicans* collected at two hours were assigned to the second cell stage and appeared to initiate the transcriptional shift in immune response earlier than in the corresponding macrophage (**Figure 4A**). In summary, the major co-stages of infection largely correspond to time of infection; however, trajectory analysis highlights an asynchronous and linear transition associated with the induction of filamentation and metabolic adaptation in *C. albicans* which, correlates with a shift from a strong to a weak pro-inflammatory gene expression profile in the host.

### Expression bimodality in host and fungal pathogen measured in single macrophages infected with live *Candida albicans*

To further examine heterogeneity in gene expression, we characterized modality of expression profiles across host-pathogen single cells, and then compared these distributions between 2 and 4 hours using a normal mixture model and Bayesian modeling framework as implemented in scDD^28^ (Methods). Overall, an average of 84.5% of genes detected across single infected macrophages displayed unimodal gene expression distributions (Figure 5A; Table S9). Highly expressed unimodal genes (top 5%) with similar expression levels at 2 and 4 hours encompassed genes involved in opsonization and *C. albicans* recognition, such as complement proteins (*C1qb, C1qc*) and galectin receptors that recognize beta-mannans (*Lgals1, Lgals3*). In phagocytosed *C. albicans*, an average of 76% had unimodal expression patterns (**Figure 5A; Table S10**). Highly expressed unimodal genes (top 5%) with similar expression levels in both co-stages were enriched in the oxidation-reduction process and defense against reactive oxygen species (enriched GO term, corrected-P < 0.05, hypergeometric distribution with Bonferroni correction; **Table S10**). These unimodal genes highlight the core genes involved in the host immune response to fungus and pathogen virulence, respectively.

**Figure 5.**
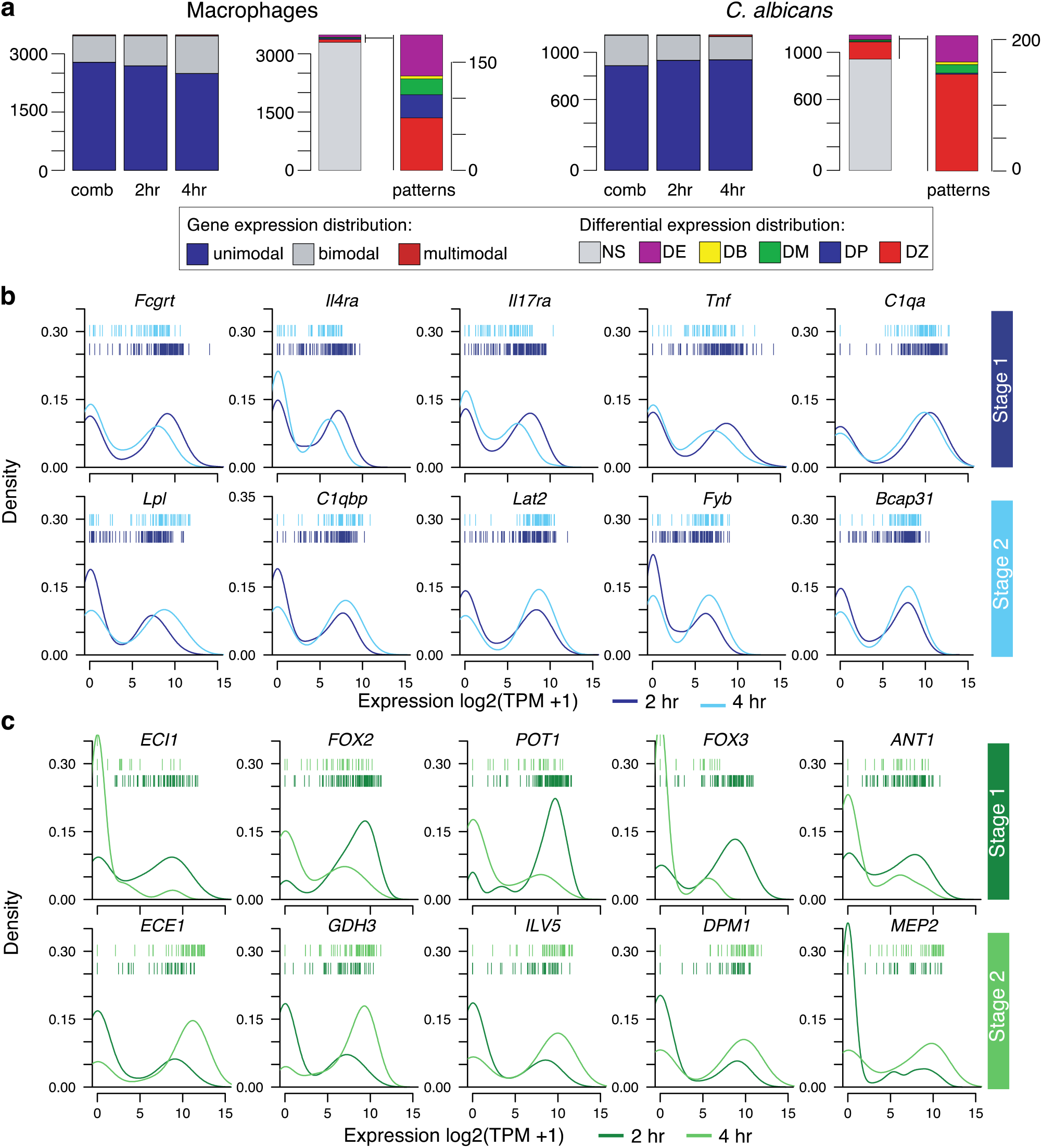
Expression variability and bimodality at the single-cell level. **(a)** Number of genes categorized as unimodal, bimodal, or multimodal (>2 components) according genes expression patterns in single, infected macrophages and corresponding phagocytized *C. albicans* across cells from 2 and 4 hours, and within each time point. Differential distributions were assigned as differential expression of unimodal genes (DE), differential modality and different component means (DB), differential modality (DM), differential proportion for bimodal genes (DP), differential proportion of zeroes (DZ), and not significant (NS). **(b)** Expression density distributions in parallel infected macrophages (*n* = 267; blue) and **(c)** phagocytized live *C. albicans* (*n* = 215; green) for 5 top marker genes in co-state1 and co-state2 across macrophages-C. *albicans* single cells at 2 and 4 hours post infection. Individual cells are plotted as bars for 2 hours (top row) and 4 hours (bottom row) for each distribution.

A subset of the genes highly expressed in co-stages of single infected macrophages and phagocytosed *C. albicans* showed bimodal expression distributions (**Figure 4B**). As bimodal transcriptional heterogeneity among single stimulated or infected cells can signify distinct immune cell expression programs^13,23^, we next examined whether subgroups of macrophages could be defined by shared bimodality of genes involved in the immune response. An average of 15% host genes showed patterns of bimodal expression among or within 2 and 4 hours (exceeded Bimodality Index (BI) threshold; Dirichlet Process Mixture of normals model; **Table S9, Figure 5A**). In addition to expression bimodality, some of these genes showed patterns of differential distributions (*e.g*. shifts in mean(s) expression, modality, and proportions of cells) across and within 2 and 4 hours as implemented in scDD package^28^. This includes genes involved in pathogen intracellular recognition and pro-inflammatory response that had a bimodal expression distribution at 2 hours but not at 4 hours (*e.g. Olr1, Tnfrsf12a*), bimodal expression only at 4 hours (*Il4ra*), or bimodal expression at 2 and 4 hours with differential mean expression (*Il21r, Il17ra; P* < 0.05, Benjamini-Hochberg adjusted Fisher’s combined test; **Figure 5B; Table S9**). As observed in macrophages infected with *Salmonella*^14^ or stimulated with LPS^14^, *Tnf* and *Il4ra* also exhibited bimodal expression patterns in single macrophages infected with *C. albicans* (**Figure 5B, top**). Additionally, we found unique subsets of genes displaying differential distributions in single macrophages infected with *C. albicans*, but not in response to bacterial stimuli, including *Il17ra* and other lectin-like receptors (*Olr1*; **Figures 5B and S12**). This suggests that variably expressed pathogen-specific receptors may play a role in these interactions, even in clonal populations of cells.

We next characterized variation in isoform usage between single macrophages during *C. albicans* infection, including immune response genes. Briefly, we calculated the frequency (percentage spliced in (PSI)) of previously annotated splicing events and identified differential isoform usage between single infected macrophages using BRIE^29^ (**Methods**). We detected differential splicing between macrophages in 144 genes, including the immune response genes *Clec4n* (*Dectin-2*), *Il10rb* and *Ifi16* (cell pairs > 2000; Bayes factor > 200; Table S10). Notably, Dectin-2 had differential exon retention between macrophage stages, with the Dectin-2 α isoform (6 exons) predominantly in stage 1, and Dectin-2 β isoform (5 exons) predominately in stage 2 (Figure 6). The truncated isoform Dectin-2 β lacks part of the intracellular domain and most of the transmembrane domain of the receptor^30^. We found that Dectin-2 β has a lower posterior probability of transmembrane helix (p = 0.80; TMHMM2) as compared as Dectin-2 α (p = 0.99; TMHMM2; Figure S13). The lack of this transmembrane region has been proposed to encode a secreted protein, which may act as an antagonist to full-length Dectin-2^30^. Activation of Dectin-2 receptors on macrophages and dendritic cells by *C. albicans* leads to Th17 T cell differentiation to assist in the immune response^31,32^. These results suggest that splicing variation among single macrophages might indicate different potentials to respond to fungal infection.

**Figure 6.**
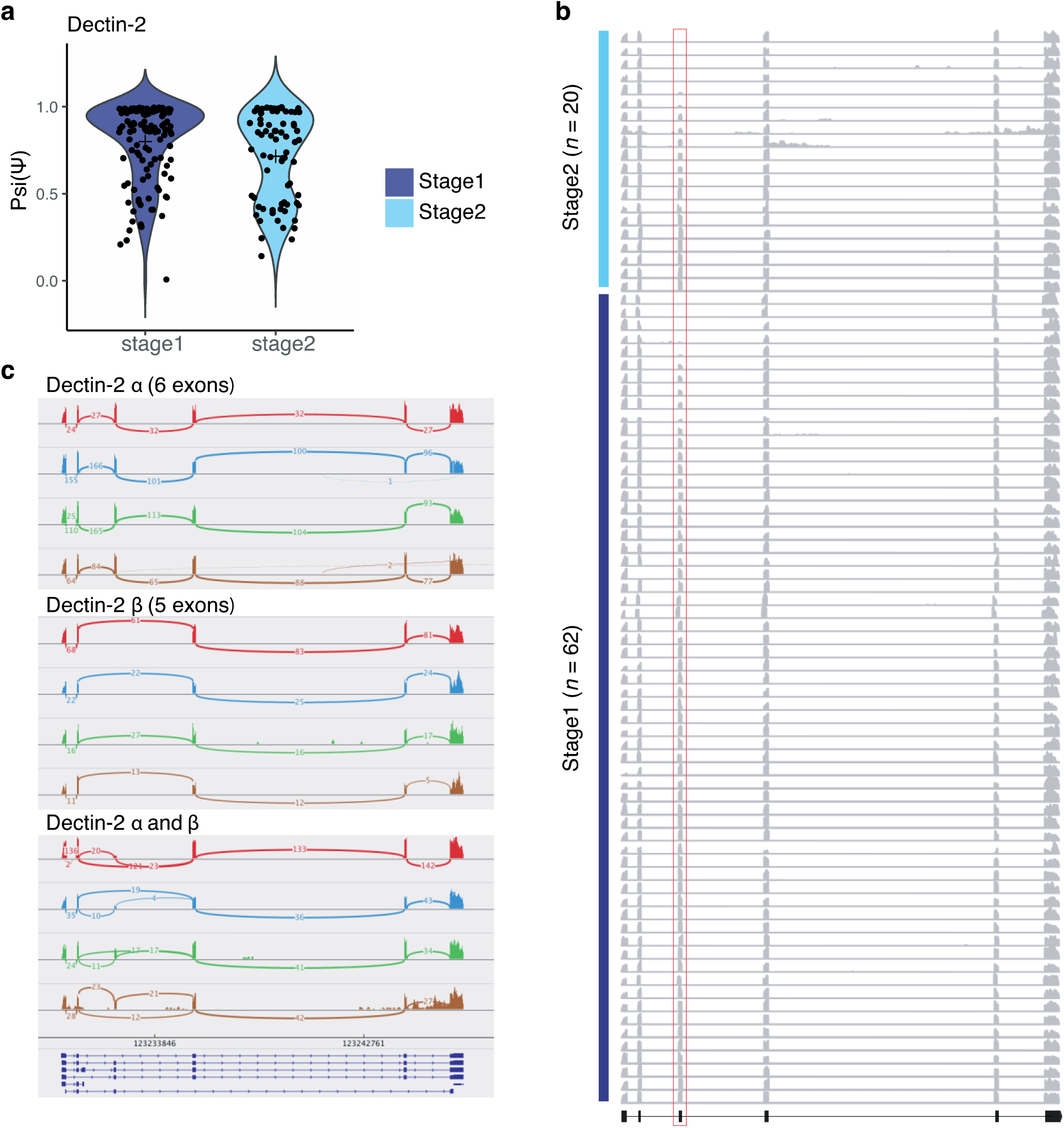
Differential splicing and isoform usage in single macrophages infected with *C. albicans*. **(a)** Differential inclusion of exon 3 alternative splicing event in Clec4n (Dectin-2) in single macrophages infected with *C. albicans*. Violin plots depict the distribution of the percent spliced-in (Psi/Ψ) scores for single macrophages in stage 1 and 2 of infection. **(b)** Shown are scRNA-seq densities for all exons of Dectin-2 across 82 single macrophages infected with *C. albicans;* 62 in stage-1 and 20 in stage-2. Each row represents a single-cell. Red box indicates the spliced exon 3 **(c)** Sashimi plots showing differential splicing between single macrophages, including the read densities and number of junction reads. Four cells were selected showing three scenarios: top: Dectin-2 α, middle Dectin-2 β, and bottom cell with both isoforms.

We next hypothesized that *C. albicans* may also demonstrate expression heterogeneity and bimodality that is linked with expression in the corresponding macrophage cell. We found an average of 23%*C. albicans* genes that showed patterns of bimodality at 2 and 4 hours, including a subset of virulence-associated genes that also showed shifts in mean expression, modality, and proportions of cells across and within 2 and 4 hours (P < 0.05, Benjamini-Hochberg adjusted Fisher’s combined test; Figure 5A; Table S10). Differential expression of those genes explained most of the variation across infection co-stages (**Figure 4A**) and were more important for *C. albicans* fate decision and trajectory (**Figure 4B**). This set of genes were enriched in cell adhesion and filamentation, oxidation-reduction process and fatty acid oxidation (enriched GO term, corrected-P < 0.05, hypergeometric distribution with Bonferroni correction; **Table S10**), including genes involved in the core filamentation network (*ALS3, ECE1, HGT2, HWP1, IHD1, OLE1*), beta-oxidation and glyoxylate cycle (*ANT1, MDH1-3, FOX2, POX1-3, PEX5, POT1*), and response to oxidative stress (*DUR1,2, GLN1, PGK1*; **Table S10; Figure 5C, bottom**). Measuring dual species gene expression in sorted infection subpopulations and in single infected cells reveals that expression heterogeneity and bimodality of genes involved in fungal morphology and adaptation are tightly co-regulated. We observed shifts in the host response during host-fungal pathogen interactions, and, in some cases, this might result in different expression levels of host immune response genes and pathogen virulence genes. This approach can further enhance our understanding of distinct infection fates and the correlated gene regulation that governs host cells and fungal pathogen interaction outcomes.

## Discussion

Host and fungal cell interactions are heterogeneous, even within clonal populations. One way to resolve this heterogeneity is to subdivide these populations by infection fate or stage to measure gene expression variability across subpopulations. Single cell approaches offer the ultimate level of subdivision to study heterogeneity and may obviate the need for sorting when run at sufficient scale. Both these approaches rely on measuring host and fungal pathogen gene expression levels using dual RNA-sequencing to provide insight as to how both species respond in each infection stage. Here, we piloted both approaches to study fungal interactions with host cells. We developed a generalizable strategy to isolate distinct host and fungal pathogen infection fates over time, including single infected cells. In both sorted populations and single infected cells, we demonstrated that gene expression changes can be measured in both host and pathogen simultaneously. This approach allowed characterization of distinct infection fates within heterogeneous host and fungal pathogen interactions. This approach could be used to better further characterize the requirement for specific host and pathogen genes for these infection responses, and single cell analysis is well suited to characterize variability in both the host and pathogen during active infections.

This dual scRNA-Seq approach builds on prior transcriptional studies of interactions between microbial pathogens and immune cells. For fungi, RNA-Seq studies have largely measured gene expression profiles of either the host or the fungal pathogen^3,6,7^ and have measured transcription profiles across infection outcomes^8–11^. To better examine heterogeneity of host-pathogen interactions, recent approaches demonstrated the use of a similar GFP live/dead reporter to measure gene expression in phagocytes infected by bacterial pathogens using scRNA-Seq^14–16^. However, these studies mainly focused on the transcriptional response of the host, as bacterial transcriptomes can be difficult to measure due to their relatively low number of transcripts^14,16^. By contrast, we have demonstrated that both host and *C. albicans* gene expression can be measured in single infected cells; further studies will be needed to examine if the same or modified approaches can be extended to other microbial-host interactions.

By examining single infected macrophages, we showed that host and pathogen exhibit transcriptional co-stages that are tightly coupled during an infection time course, providing a high-resolution view of host-fungal interactions. This also revealed that expression heterogeneity of key genes in both infected macrophages and in phagocytosed *C. albicans* may contribute to infection outcomes. We identified two, time dependent linear co-stages of host-fungal pathogen interaction, with potential intermediate stages in some single cells suggested by pseudo-time analysis. The initial co-stage is characterized by induction of a pro-inflammatory host profile after 2 hours of interaction with *C. albicans* that then decreased by 4 hours. This is consistent with studies in human macrophages, where pro-inflammatory macrophages that interact with *C. albicans* for 8 hours or longer skew toward an anti-inflammatory proteomic profile^33^. Both commensal and invasive stages of *C. albicans* infection are impacted by the balance between pro-inflammatory and anti-inflammatory responses^34^. In single macrophages infected with *C. albicans*, the shift to an anti-inflammatory state, including upregulation of genes involved in the activation of inflammasomes, was coupled with the activation of filamentation and cell-wall remodeling in *C. albicans*. Previous work has shown that *C. albicans* can switch from yeast to hyphal growth within the nutrient-deprived, acidic phagosome^35^ and can escape by rupturing the macrophage membrane during intra-phagocytic hyphal growth^36^. Other mechanisms of escape include the activation of macrophage programmed cell death pathways, including the formation of inflammasomes and pyroptosis^37^, or cell damage induced by a cytolytic peptide toxin (Candidalysin *ECE1*)^38^. Our analysis of single cells revealed bimodal expression in genes involved in these processes, and time analysis suggests that an initial shift in expression of filamentation and cell-wall remodeling programs in phagocytosed *C. albicans* rapidly result in down-regulation of the pro-inflammatory state of the host cells. This supports the hypothesis that an asynchronous *C. albicans* yeast-to-hyphae transition within the macrophage could drive expression heterogeneity in the host. While gene expression patterns are tightly correlated in host and fungal pathogen and the progression in co-stages of infection appears largely linear, greater time resolution across more diverse cell populations will be needed to fully resolve the expression heterogeneity among individual infected macrophages and phagocytosed *C. albicans*. Further work using this approach could also examine the role of specific genes in both *C. albicans* and the host to clarify the drivers of transcriptional responses and heterogeneity in host-fungal pathogen infection fates.

We found that both single infected macrophages and the corresponding phagocytosed *C. albicans* cells exhibit expression bimodality for a subset of genes. The expression bimodality observed for the host as well as the pathogen is consistent with the evolutionary concept of bet hedging^39^. Both cell types may rely on stochastic diversification of phenotypes to improve their survival rate in the event of an encounter with the other cell type. For instance, clonal populations of *C. albicans* that find themselves in the unpredictable, changing environment of a host phagocyte could increase the chance of survival by varying the expression of key genes involved in the response. A similar scenario may provide an advantage for phagocytes that encounter pleomorphic *C. albicans*. These strategies are noted to occur within microbial populations, where a small fraction of “persister” cells might be capable of surviving exposure to lethal doses of antimicrobial drugs as a bet-hedging strategy^40^. Larger numbers of host-pathogen pairs collected over a longer time course could disambiguate how expression co-stages in *C. albicans* and macrophages results in distinct infection fates.

Heterogenous transcriptional responses are important to consider in the treatment of fungal infections. Genes expressed uniformly among fungal cells in a population may be more effective therapeutic targets than the products of genes expressed by only a subset of cells. A comparison of transcriptional signal of drug-treated single infected cells to non-treated cells would determine if all phagocytosed *C. albicans* cells co-regulate uniformly regulate genes involved in the response to drug treatment or if there is evidence of bimodal expression for genes involved in drug targets or efflux pumps, for example. Additionally, single cell analysis revealed bimodal expression and differential splicing of *Clec4n (Dectin-2),a* C-type lectin receptor that recognizes diverse fungal pathogens, as well as some bacteria and parasites^41^. The switch from producing a full length isoform to one missing an exon involved in signaling is related to the shift to an anti-inflammatory state. Parallel host-fungal pathogen expression profiling at single-cell level could not only allow researchers to pinpoint changes in immune recognition and response pathways but also measure the dynamics of other stimuli, for example to characterize how new drug treatments affect both pathogen and host cells. While we used sorting to enrich for cell populations of interest, as scRNA-Seq microfluidic platforms continue to develop and profiling thousands of single cells becomes more cost effective, it will become tractable to interrogate more complex samples, including those containing multiple host cell types and non-clonal pathogens. These data will provide fine mapping of diverse microbial immune responses and communication between immune cells. Importantly, this approach will allow further investigation into how fungal phenotypic and expression heterogeneity drives host responses and provide a systems view of these interactions.

## Methods

### *Candida albicans* reporter strain construction

The reporter construct used in this study was prepared by integrating the GFP and mCherry fluorescent tags driven by the bi-directional *ADH1* promoter and a nourseothricin resistance (NAT^R^) cassette at the Neut5L locus of *Candida albicans* strain CAI4-F2^42^. CAI4-F2 harbors a homozygous *URA3* deletion which makes it less filamentous than other common laboratory strains of *C. albicans*, including SC5314 (**Figure S14**); this allowed more *C. albicans* cells to be sorted (see **Methods; Macrophage and *Candida albicans* infection assay**). Briefly, the pUC57 vector containing mCherry driven by *ADH1* promoter (Bio Basic) was digested and this portion of the plasmid was ligated into a pDUP3 vector^43^ containing GFP, also driven by *ADH1* promoter, a NAT^R^ marker, and homology to the Neut5L locus. The resulting plasmid was linearized and introduced via homologous recombination into a neutral genomic locus, Neut5L, using chemical transformation protocol with lithium acetate. Transformation was confirmed via colony PCR and whole-genome sequencing. A whole genome library was created from strain CAI4-F2-Neut5L-NAT1-mCherry-GFP using Nextera-XT library construction strategy(Illumina) and sequenced on an Illumina MiSeq (150×150 paired end sequencing; approximately 28 million reads). Sequencing reads were *de novo* assembled using dipSPAdes^44^. Scaffolds were queried back to the plasmid sequence used to transform CAI4-F2 using BLAST^45^. Sequencing coverage was visualized using IGV^46^.

### Macrophage and *Candida albicans* infection assay

Primary, bone derived macrophages (BMDMs) were derived from bone marrow cells collected from the femur and tibia of C57BL/6, female mice. All mouse work was performed in accordance with the Institutional Animal Care and Use Committees (IACUC) and with relevant guidelines at the Broad Institute and Massachusetts Institute of Technology, with protocol 0615-058-1. Primary bone marrow cells were grown in “C10” media as previously described^47^ and supplemented with macrophage colony stimulating factor (M-CSF) (ThermoFisher Scientific) at final concentration of 10 ng/ml, to promote differentiation into macrophages. C10 media was not supplemented with uridine; this led to reduced CAI4-F2 filamentation (compared to wild type SC5314) and allowed more *C. albicans* cells to be sorted. Cultures were then stained with F4/80 (Biolegend) to ensure that ~95% of the culture had differentiated into macrophages. For the infection RNA-sequencing experiment, BMDMs were seeded in 6 well plates (Falcon). Two days prior to the start of the infection experiment, yeast strains were revived on rich media plates. One day prior to the infection experiment, yeast were grown in 3 ml overnight cultures in rich media at 30 °C. On the day of the infection experiment, macrophages were stained with CellMask Deep Red plasma membrane stain (diluted 1:1000) (ThermoFisher Scientific). Macrophages and stain were incubated at 37 °C for 10 minutes, then macrophages were washed twice in 1X PBS. 2 hours prior to infection, yeast cells were acclimated to macrophage media (RPMI 1640 no phenol red, plus glutamine, ThermoFisher Scientific) at 37 °C prior to the infection. Yeast cells were then counted and seeded in a ratio of 1 *C. albicans* cell to 2 macrophage cells. Yeast and macrophages were then co-incubated at 37 °C (5% C02). At the indicated time point, media was removed via aspiration, 1 ml of 1X TrypLE, no phenol red (ThermoFisher Scientific) was added to each well and incubated for 10 minutes. After vigorous manual pipetting, 2 wells for each time point were combined into one tube. Each time point was run in biological triplicate. Samples were then spun down at 37 °C, 300g for 10 minutes and resuspended in 1 ml PBS + 2% FCS and placed on ice until FACS. RNAlater was used shortly after sorting, as this reagent leads to decreased GFP expression^49^. To control for gene expression changes that may be induced by FACS, only sorted samples were used in comparative analyses.

### Fluorescence-activated cell sorting (FACS)

Samples were sorted on the BDSORP FACSAria running the BD FACSDIVA8.0 software into 1X PBS and then frozen at −80 °C until RNA extraction. Cells were sorted into the following sub populations: (i) macrophages infected with live *C. albicans* (GFP+, mCherry+, Deep red+), (ii) macrophages infected with dead, phagocytosed *C. albicans* (GFP-, mCherry+, Deep red+), (iii) macrophages exposed to *C. albicans* (GFP-, mCherry-, Deep red+) and (iv) *C. albicans* exposed to macrophages (GFP+, mCherry+, Deep red-) Single cells were sorted into 5 ul of RLT 1%β-Mercaptoethanol in a 96 well plate (Eppendorf) and frozen at −80°C.

### RNA extraction, evaluation of RNA quality

RNA was extracted from population samples using the Qiagen RNeasy mini kit. All samples were resuspended in RLT (Qiagen) + 1%β-Mercaptoethanol (Sigma) and subjected to 3 minutes of bead beating with .5mm zirconia glass beads (BioSpec Products) in a bead mill. Single macrophages infected with *C. albicans* were directly with dead, phagocytosed by sorting cells into a 96 well plate containing 5 ul of RLT (Qiagen) + 1%β-Mercaptoethanol (Sigma).

### cDNA synthesis and library generation

For population samples, the RT reaction was carried out with the following program, as described^50^, with the addition of RNAse inhibitor (ThermoFisher) at 40U/ul. cDNA was generated from single cells based on the Smart-seq2 method as described previously^51^, with the addition of RNase inhibitor was used at 40 U/ul (ThermoFisher) and 3.4 ul of 1 M trehalose was added to the RT reaction. All libraries were constructed using the Nextera XT DNA Sample Kit (Illumina) with custom indexed primers as described^51^. Infection subpopulation samples were sequenced on an Illumina NextSeq (37 x 38 cycles). *Candida* only samples were sequenced on an Illumina MiSeq (75 x 75 cycles). Single infected cells were sequenced on Illumina’s NextSeq (75×75 cycles).

### Read processing and transcript quantification of population-RNA-Seq

Basic quality assessment of Illumina reads and sample demultiplexing was done with Picard version 1.107 and Trimmomatic^52^. Samples profiling exclusively the mouse transcriptional response were aligned to the mouse transcriptome generated from the v. Dec. 2011 GRCm38/mm10 and a collection of mouse rRNA sequences from the UCSC genome website^53^. Samples profiling exclusively the yeast transcriptional response were aligned to transcripts from the *Candida albicans* reference genome SC5314 version A21-s02-m09-r10 downloaded from the *Candida* Genome Database (http://www.candidagenome.org).

Samples from the infection assay profiling in parallel host and fungal transcriptomes, were aligned to a “composite transcriptome” made by combining the mouse transcriptome described above and the *C. albicans* transcriptome described above. To evaluate read mappings, BWA aln (BWA version 0.7.10-r789, http://bio-bwa.sourceforge.net/)^54^ was used to align reads, and the ‘XA tag’ was used for read enumeration and separation of host and pathogen sequenced reads. Multi-reads (reads that aligned to both host and pathogen transcripts) were discarded, representing only an average of 2.6% of the sequenced reads. Then, each host or pathogen sample file were aligned to its corresponding reference using Bowtie2^55^ and RSEM (RNA-Seq by expectation maximization; v.1.2.21). Transcript abundance was estimated using transcripts per million (TPM). Since parallel sequencing of host and pathogen from single macrophages increased the complexity of transcripts measured compared to studies of only host cells alone, we detected lower number of transcripts of macrophages as compared with other studies using phagocytes and similar scRNA-seq methods^14,15,23^.

### Differential gene expression analysis of population-RNA-Seq

TMM-normalized ‘transcripts per million transcripts’ (TPM) for each transcript were calculated, and differentially expressed transcripts were identified using edgeR^56^, all as implemented in the Trinity package version 2.1.1^57^. Genes were considered differentially expressed if they had a 4-fold change difference (> 4 FC) in TPM values and a false discovery rate below or equal to 0.001 (FDR < 0.001), unless specified otherwise.

### Read processing and transcript quantification of single-cell RNA-Seq

BAM files were converted to merged, demultiplexed FASTQ format using the Illumina Bcl2Fastq software package v2.17.1.14. For the RNA-Seq of sorted population samples, paired-end reads were mapped to the mouse transcriptome (GRCm38/mm10) or to the *Candida albicans* transcriptome strain SC5314 version A21-s02-m09-r10 using Bowtie2^55^ and RSEM (RNA-Seq by expectation maximization; v.1.2.21)^58^. Transcript abundance was estimated using transcripts per million (TPM). For read mapping counts paired-reads were aligned to the ‘composite reference’ as described above.

For each single macrophage infected with *C. albicans*, we quantified the number of genes for which at least one read was mapped (TPM > 1). We filtered out low-quality macrophage or *C. albicans* cells from our data set based on a threshold for the number of genes detected (a minimum of 2,000 unique genes per cell for macrophages, and 600 unique genes per cell for *C. albicans*, and focused on those single infected macrophages that have good number of transcript detected in both host and pathogen (**Figure S9A**). For a given sample, we define the filtered gene set as the genes that have an expression level exceeding 10 TPM in at least 20% of the cells. After cell and gene filtering procedures, the expression matrix included 3,254 transcripts for the macrophages and 915 transcripts for *C. albicans*. To estimate the number of *C. albicans* in each macrophage, we measured the correlation of GFP levels from FACs with the total number of transcripts detected in live, phagocytosed *C. albicans* cells (at least 1 TPM) but found only a modest correlation between these two metrics (R^2^ = 0.52).

To eliminate the non-biological associations of the samples based on plate-based processing and amplification, single-cell expression matrices were log-transformed (log(TPM + 1)) for all downstream analyses, most of which were performed using the R software package Seurat (https://github.com/satijalab/seurat). In addition, we do not find substantial differences in the number of sequenced reads and detected genes between samples. We separately analyzed two comparisons of macrophages-*C. albicans* cells: i) single macrophages infected - live phagocytosed *C. albicans* cells at 2 and 4 hours (macrophages = 267; *C. albicans* cells = 215; macrophage-C. *albicans* = 156); and *ii*) macrophages infected and live or dead, phagocytosed *C. albicans* cells at 4 hours (macrophages = 142; *C. albicans* cells = 71). These numbers of macrophages and *C. albicans* cells are the total that met the described QC filters.

### Detection of variation across single, infected cells

To examine if cell to cell variability existed across a wide range of population expression levels, we analyzed the variation and the intensity of non-unimodal distribution for each gene across single macrophages and *C. albicans* cells. Briefly, we determined the distribution of the average expression (μ), and the dispersion of expression (σ^2^; normalized coefficient of variation), placing each gene into bins, and then calculating a z-score for dispersion within each bin to control for the relationship between variability and average expression as implemented R package Seurat^25^.

### Detection of variable genes, cell clustering and trajectory

To classify the single cell RNA-Seq from macrophages and *C. albicans*, the R package Seurat was used^25^. We first selected variable genes by fitting a generalized linear model to the relationship between the squared co-efficient of variation (CV) and the mean expression level in log/log space, and selecting genes that significantly deviated (p-value < 0.05) from the fitted curve, as implemented in Seurat, as previously described^15^. Then highly variable genes (CV > 1.25; p-value < 0.05) were used for principle component analysis (PCA), and statistically significant determined for each PC using JackStraw plot function. Significant PCs (p-value < 0.05) were used for two-dimension t-distributed stochastic neighbor embedding (tSNE) to define subgroups of cells we denominated host-pathogen co-states. We identified differentially expressed genes (corrected-p < 0.05) between co-states using a likelihood-ratio test (LRT) for single-cell differential expression^26^ as implemented in Seurat. To perform pseudo-time analysis we Monocle 2.8.0 method^27^ using filtered normalized genes from Seurat. We reordered cells in pseudo-time along a trajectory using the top 50 differentially expressed genes, and identified genes with most significant changes as a function of progress along the trajectory using the branched expression analysis modeling (BEAM; q-value < 1e-4).

### Detection of differential expression distributions and bimodality

To detect which genes have different expression distributions single infected macrophage and live *C. albicans* we compared the distributions of gene expression within single infected macrophage and live *C. albicans*, across and between 2 and 4 hours, and identified genes showing evidence of differential distribution using a Bayesian modeling framework as implemented in scDD^28^. We used the permutation test of the Bayes Factor for independence of condition membership with clustering (n permutations = 1000), and test for test for a difference in the proportion of zeroes (testZeroes=TRUE). A gene was considered differentially distributed using Benjamini-Hochberg adjusted p-values test (p-value < 0.05).

### Detection of differential splicing

To detect alternative splicing and determine variation in isoform usage between single macrophages during *C. albicans* infection we calculated exon splicing rates in individual macrophages using the data-dependent module of BRIE v0.2.0^29^. BRIE calls splicing at predefined cassette exons and quantifies splicing using exon reads in single-cell data. We used the processed annotation file and sequence features for mouse (mm10 GRCm38.p5, ANNO_FILE = SE.gold, FACTOR_FILE = mouse_factors.SE.gold). Differential splicing was performed using the brie-diff module for all pairwise comparisons, and genes with differential splicing between single macrophages were defined as cell pairs > 2000 and Bayes factor > 200.

### Functional biological enrichment analyses

For *C. albicans*, Gene ontology (GO) term analysis was performed in through the *Candida* Genome Database GO Term Finder and GO Slim Mapper (http://www.candidagenome.org^59^). GO terms were considered significantly enriched in a cluster or set of genes if we found a GO term corrected *p-value* lower than 0.05 using hypergeometric distribution with Bonferroni correction

For macrophages, Ingenuity Pathway analysis (IPA) was performed. We investigated biological relationships, canonical pathways and Upstream Regulator analyses as part of IPA software. This allowed us to assess the overlap between significantly DEGs and an extensively curated database of target genes for each of several hundred known regulatory proteins. Clusters or set of genes were considered significantly enriched if we found enriched a *-log*(*p-value*) greater than 1.3 (*i.e*. p-value < 0.05) and z-score greater than 2 as recommended by the IPA software.

### Data and biological materials availability statement

All sequence data for this project has been deposited in the SRA under Bioproject PRJNA437988. Raw and processed data for gene expression analysis was deposited in the GEO under GSE111731. Whole genome sequence data of CAI4-F2-Neut5L-NAT1-mCherry-GFP has been deposited at the NCBI SRA under SRX4924342. The CAI4-F2-Neut5L-NAT1-mCherry-GFP reporter strain is available upon request to RPR or from the ATCC BEI (accession pending).

## Acknowledgments

We thank Aviv Regev and members of her lab for providing support for all the experimental work in this paper. We also thank Raktima Raychowdhury for help with preparation of the primary BMDM cells and Anh Hoang, Mehment Toner, and Daniel Irima for providing microwells used in preliminary experiments. This project has been funded in whole or in part with Federal funds from the National Institute of Allergy and Infectious Diseases, National Institutes of Health, Department of Health and Human Services, under award N°: U19AI110818. CBF was supported by a Helen Hay Whitney postdoctoral fellowship, TD was supported by NIAID and WPI.

## Author Contributions

CBF, DAT, RPR, and CAC designed the study. TD, CBF and BYL carried out experiments. JFM analyzed the data and prepared figures and tables. JFM, TD, RPR and CAC wrote the initial draft of the manuscript, which was revised with input from all authors. All authors read and approved final manuscript.

## References

1. Brown, G. D. Innate antifungal immunity: the key role of phagocytes. Annu. Rev. Immunol. 29, 1–21 (2011).

2. Huffnagle, G. B. & Noverr, M. C. The emerging world of the fungal microbiome. Trends Microbiol. 21, 334–341 (2013).

3. Lorenz, M. C., Bender, J. A. & Fink, G. R. Transcriptional response of *Candida albicans* upon internalization by macrophages. Eukaryot. Cell 3, 1076–1087 (2004).

4. Frohner, I. E., Bourgeois, C., Yatsyk, K., Majer, O. & Kuchler, K. *Candida albicans* cell surface superoxide dismutases degrade host-derived reactive oxygen species to escape innate immune surveillance. Mol. Microbiol. 71, 240–252 (2009).

5. Seider, K., Heyken, A., Lüttich, A., Miramón, P. & Hube, B. Interaction of pathogenic yeasts with phagocytes: survival, persistence and escape. Curr. Opin. Microbiol. 13, 392–400 (2010).

6. Smeekens, S. P. et al. Functional genomics identifies type i interferon pathway as central for host defense against *Candida albicans*. Nat. Commun. 4, (2013).

7. Quintin, J. et al. *Candida albicans* infection affords protection against reinfection via functional reprogramming of monocytes. Cell Host Microbe 12, 223–232 (2012).

8. Hebecker, B. et al. Dual-species transcriptional profiling during systemic candidiasis reveals organ-specific host-pathogen interactions. Sci Rep 6, 36055 (2016).

9. Liu, Y. et al. New signaling pathways govern the host response to *C. albicans* infection in various niches. Genome Res. 125, 679–689 (2015).

10. Niemiec, M. J. et al. Dual transcriptome of the immediate neutrophil and *Candida albicans* interplay. BMC Genomics 18, 696 (2017).

11. Tierney, L. et al. An interspecies regulatory network inferred from simultaneous RNA-seq of *Candida albicans* invading innate immune cells. Front. Microbiol. 3, 1–14 (2012).

12. Erwig, L. P. & Gow, N. a. R. Interactions of fungal pathogens with phagocytes. Nat. Rev. Microbiol. 14, 163–176 (2016).

13. Shalek, A. K. et al. Single-cell transcriptomics reveals bimodality in expression and splicing in immune cells. Nature 498, 236–40 (2013).

14. Avraham, R. et al. Pathogen Cell-to-Cell Variability Drives Heterogeneity in Host Immune Responses. Cell 162, 1309–1321 (2015).

15. Saliba, A.-E. E. et al. Single-cell RNA-seq ties macrophage polarization to growth rate of intracellular Salmonella. Nat. Microbiol. 2, 1–8 (2016).

16. Avital, G. et al. scDual-Seq: mapping the gene regulatory program of *Salmonella* infection by host and pathogen single-cell RNA-sequencing. Genome Biol. 18, 200 (2017).

17. Martin, R. et al. A core filamentation response network in *Candida albicans* is restricted to eight genes. PLoS One 8, e58613 (2013).

18. Issi, L. et al. Zinc cluster transcription factors alter virulence in *Candida albicans*. Genetics 205, 559–576 (2017).

19. Du, H. et al. Roles of *Candida albicans* Gat2, a GATA-Type Zinc Finger Transcription Factor, in Biofilm Formation, Filamentous Growth and Virulence. PLoS One 7, e29707 (2012).

20. Bruno, V. M. et al. Transcriptomic Analysis of Vulvovaginal Candidiasis Identifies a Role for the NLRP3 Inflammasome. MBio 6, e00182–15 (2015).

21. Lionakis, M. S. et al. CX3CR1-dependent renal macrophage survival promotes *Candida* control and host survival. J. Clin. Invest. 123, 5035–51 (2013).

22. Leonardi, I. et al. CX3CR1+mononuclear phagocytes control immunity to intestinal fungi. Science 359, 232–236 (2018).

23. Shalek, A. K. et al. Single-cell RNA-seq reveals dynamic paracrine control of cellular variation. Nature 510, 363–9 (2014).

24. Van Der Maaten, L. J. P. & Hinton, G. E. Visualizing high-dimensional data using t-sne. J. Mach. Learn. Res. 9, 2579–2605 (2008).

25. Macosko, E. Z. et al. Highly Parallel Genome-wide Expression Profiling of Individual Cells Using Nanoliter Droplets. Cell 161, 1202–1214 (2015).

26. McDavid, A. et al. Data exploration, quality control and testing in single-cell qPCR-based gene expression experiments. Bioinformatics 29, 461–467 (2013).

27. Trapnell, C. et al. The dynamics and regulators of cell fate decisions are revealed by pseudotemporal ordering of single cells. Nat. Biotechnol. 32, 381–386 (2014).

28. Korthauer, K. D. et al. A statistical approach for identifying differential distributions in single-cell RNA-seq experiments. Genome Biol. 17, 222 (2016).

29. Huang, Y. & Sanguinetti, G. BRIE: transcriptome-wide splicing quantification in single cells. Genome Biol. 18, 123 (2017).

30. Gavino, A. C. P., Chung, J.-S., Sato, K., Ariizumi, K. & Cruz, P. D. Identification and expression profiling of a human C-type lectin, structurally homologous to mouse dectin-2. Exp. Dermatol. 14, 281–288 (2005).

31. Saijo, S. et al. Dectin-2 recognition of α-mannans and induction of Th17 cell differentiation is essential for host defense against *andida albicans*. Immunity 32, 681–691 (2010).

32. Robinson, M. J. et al. Dectin-2 is a Syk-coupled pattern recognition receptor crucial for Th17 responses to fungal infection. J. Exp. Med. 206, 2037–2051 (2009).

33. Reales-Calderón, J. A., Aguilera-Montilla, N., Corbí, Á. L., Molero, G. & Gil, C. Proteomic characterization of human proinflammatory M1 and anti-inflammatory M2 macrophages and their response to *Candida albicans*. Proteomics 14, 1503–1518 (2014).

34. Jouault, T. et al. Host responses to a versatile commensal: PAMPs and PRRs interplay leading to tolerance or infection by *Candida albicans*. Cell. Microbiol. 11, 1007–15 (2009).

35. Westman, J., Moran, G., Mogavero, S., Hube, B. & Grinstein, S. *Candida albicans* Hyphal Expansion Causes Phagosomal Membrane Damage and Luminal Alkalinization. MBio 9, e01226–18 (2018).

36. Bain, J. M. et al. *Candida albicans* Hypha Formation and Mannan Masking of β-Glucan Inhibit Macrophage Phagosome Maturation. MBio 5, e01874–14 (2014).

37. Wellington, M., Koselny, K., Sutterwala, F. S. & Krysan, D. J. *Candida albicans* triggers NLRP3-mediated pyroptosis in macrophages. Eukaryot. Cell 13, 329–340 (2014).

38. Kasper, L. et al. The fungal peptide toxin Candidalysin activates the NLRP3 inflammasome and causes cytolysis in mononuclear phagocytes. Nat. Commun. 9, 4260 (2018).

39. de Jong, I. G., Haccou, P. & Kuipers, O. P. Bet hedging or not? A guide to proper classification of microbial survival strategies. BioEssays 33, 215–223 (2011).

40. Lewis, K. Persister Cells. Annu. Rev. Microbiol. 64, 357–372 (2010).

41. Kerscher, B., Willment, J. A. & Brown, G. D. The Dectin-2 family of C-type lectin-like receptors: An update. Int. Immunol. 25, 271–277 (2013).

42. Selmecki, A., Bergmann, S. & Berman, J. Comparative genome hybridization reveals widespread aneuploidy in *Candida albicans* laboratory strains. Mol. Microbiol. 55, 1553–1565 (2005).

43. Gerami-Nejad, M., Zacchi, L. F., McClellan, M., Matter, K. & Berman, J. Shuttle vectors for facile gap repair cloning and integration into a neutral locus in *Candida albicans*. Microbiol. (United Kingdom) 159, 565–579 (2013).

44. Safonova, Y., Bankevich, A. & Pevzner, P. A. in Journal of Computational Biology 22, 265–279 (Springer, Cham, 2014).

45. Altschul, S. F. et al. Gapped BLAST and PSI-BLAST: a new generation of protein database search programs. Nucleic Acids Res. 25, 3389–402 (1997).

46. Thorvaldsdottir, H. et al. Integrative Genomics Viewer (IGV): high-performance genomics data visualization and exploration. Brief. Bioinform. 14, 178–92 (2013).

47. Stubbs, M. et al. MLL-AF9 and FLT3 cooperation in acute myelogenous leukemia: development of a model for rapid therapeutic assessment. Leukemia 22, 66–77 (2008).

48. Schindelin, J. et al. Fiji: an open-source platform for biological-image analysis. Nat. Methods 9, 676–682 (2012).

49. Zaitoun, I., Erickson, C. S., Schell, K. & Epstein, M. L. Use of RNAlater in fluorescence-activated cell sorting (FACS) reduces the fluorescence from GFP but not from DsRed. BMC Res. Notes 3, 328 (2010).

50. Satija, R., Farrell, J. A., Gennert, D., Schier, A. F. & Regev, A. Spatial reconstruction of single-cell gene expression data. 33, (2015).

51. Villani, A.-C. et al. Single-cell RNA-seq reveals new types of human blood dendritic cells, monocytes, and progenitors. Science (80-.). 356, (2017).

52. Bolger, A. M., Lohse, M. & Usadel, B. Trimmomatic: a flexible trimmer for Illumina sequence data. Bioinformatics 30, 2114–2120 (2014).

53. Karolchik, D. et al. The UCSC Table Browser data retrieval tool. Nucleic Acids Res. 32, 493D–496 (2004).

54. Li, H. & Durbin, R. Fast and accurate long-read alignment with Burrows-Wheeler transform. Bioinformatics 26, 589–95 (2010).

55. Langmead, B. & Salzberg, S. L. Fast gapped-read alignment with Bowtie 2. Nat. Methods 9, 357–359 (2012).

56. Robinson, M. D., McCarthy, D. J. & Smyth, G. K. edgeR: a Bioconductor package for differential expression analysis of digital gene expression data. Bioinformatics 26, 139–40 (2010).

57. Haas, B. J. et al. De novo transcript sequence reconstruction from RNA-seq using the Trinity platform for reference generation and analysis. Nat. Protoc. 8, 1494–512 (2013).

58. Li, B. & Dewey, C. N. RSEM: accurate transcript quantification from RNA-Seq data with or without a reference genome. BMC Bioinformatics 12, 323 (2011).

59. Inglis, D. O. et al. The *Candida* genome database incorporates multiple *Candida* species: multispecies search and analysis tools with curated gene and protein information for *Candida albicans* and *Candida glabrata*. Nucleic Acids Res. 40, D667–D674 (2012).

