## Supplementary Information for "Coordinated host-pathogen transcriptional dynamics revealed using sorted subpopulations and single, *Candida albicans* infected macrophages"

**Supplementary Notes**

1. **Characterization of heterogeneous infection subpopulations in *ex vivo* macrophage and *Candida albicans* interactions**

The number of RNA-Seq reads and transcripts detected for both host and fungal pathogen subpopulations was sufficient for differential expression analysis in most samples and to widely profile parallel transcriptional responses. We aligned reads to a composite reference of both mouse and *C. albicans* transcriptomes (**Methods**) and found that the fraction of mapped reads for host and pathogen was correlated with the percent of sorted cells for each subpopulation (**Figures 1B**, **S2B**; **Table S1**). In subpopulations containing both host and pathogen, the fraction of reads averaged 87% macrophage and 13% *C. albicans* for macrophages infected with live fungal cells, and 95% macrophage and 5% *C. albicans* for macrophages containing dead fungal cells. For the subpopulation of macrophages infected with live fungal cells, between 1.4 and 34.0 million reads mapped to host transcriptome, while between 0.3 to 13.3 million reads mapped to *C. albicans* transcriptome. *Candida* reads in this subpopulation increased over the time course (0,1, 2 and 4 hours; **Figure 1B)**, reflecting ongoing engulfment and/or *C. albicans* division within the macrophage. In subpopulations of dead phagocytosed cells, read counts were lower than other subpopulations and counts increased over time, reflecting their smaller proportion of sorted cells that also increased over time (**Figure S2B**). In subpopulations of macrophages infected with live fungus, an average of 10,333 host and 4,567 *C. albicans* genes were detected (at least 1 fragment per replicate across all samples; **Methods**; **Figure S3A**; **Table S1**). Fewer transcripts were detected in subpopulations of macrophages infected with dead *C. albicans* (an average of 3,214 host transcripts and 983 fungal transcripts **Figure S3A**) and had modestly correlated biological replicates (*e.g.* Pearson’s *r* < 0.56). Lower coverage was expected for this subpopulation, as only up to 3% of each sample collected at each time point was comprised of macrophages infected with dead *C. albicans* (**Figure S2B**). Therefore, we primarily focused the differential expression analysis on subpopulations of macrophages or *C. albicans* exposed, and macrophages infected with live *C. albicans*, which had high transcriptome coverage, and highly correlated biological replicates (*e.g.* Pearson’s *r* 0.96 and 0.92 in macrophages and *Candida* at 4 hours, respectively; **Figure S3B**). These results indicate that we have established a robust system for measuring host and fungal pathogen transcriptional signal during phagocytosis in sorted infection subpopulations.

2. **Subpopulations of phagocytosed *C. albicans* adapt to macrophages by switching metabolic pathways and regulating cell morphology**

We next examined how *C. albicans* gene expression varied across unexposed, exposed and phagocytosed cells over time. Using *k-means* clustering, we identified sets of genes with similar expression patterns; the major patterns of expression across time were either induced (cluster 1, 2 and 3) or repressed (cluster 4 and 5) in the live, phagocytosed subpopulation relative to all other *C. albicans* infection fates (**Figures 2B**, **S4A**). A large number of genes were differentially expressed in phagocytosed *C. albicans* (732 genes relative to unexposed across all time points) compared with the number of DEGs found in exposed but un-engulfed *C. albicans* (82 genes relative to unexposed across all time points). Comparing the phagocytosed and un-engulfed *C. albicans* subpopulations at each time point, the major differential response was found at 1 hour, highlighting a rapid and specific transcriptional response upon macrophage phagocytosis. Many of these genes maintained high expression levels throughout the 4-hour infection time course (**Table S2**; **Figures 2B**, **2E**).

**Supplementary Figures**

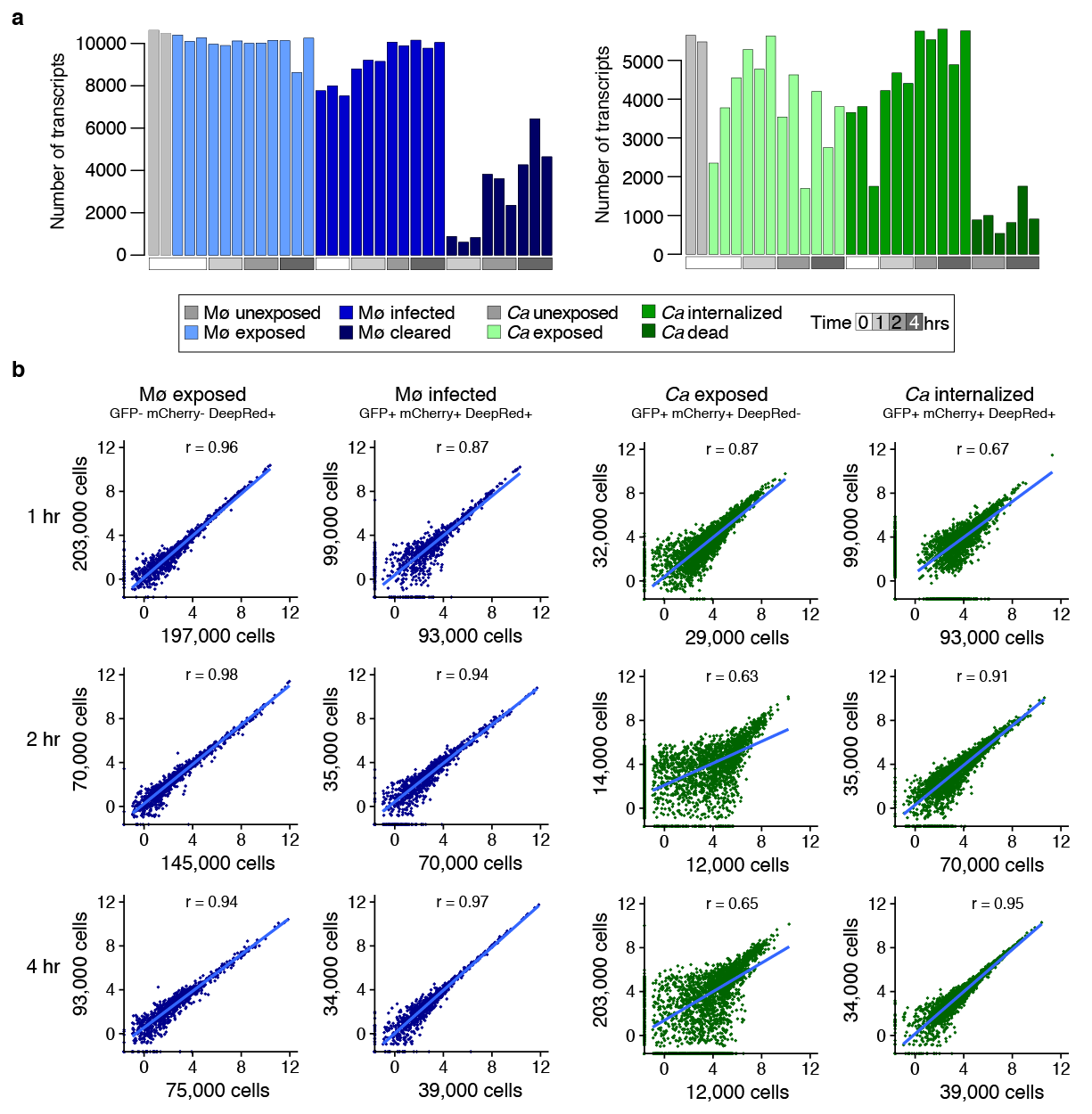

**Figure S1. Parallel captured transcripts coverage and sample correlation in subpopulation of *Candida albicans*-macrophages infection outcome. (a)** Number of transcripts simultaneously detected (TPM >1) in distinct subpopulations of macrophages (Mø; left; orange) and *C. albicans* (*Ca*; right; green). **(b)** Analysis of the correlation (Pearson correlation coefficient) of gene expression for sorted subpopulations replicates during macrophage-*C.albicans* interaction. The axes indicate number of sorted cell per replicate. Three subpopulations are depicted: macrophages infected-live *C. albicans* (GPF+, mCherry+, DeepRed+), macrophages exposed (GPF-, mCherry-, DeepRed+), and *C. albicans* exposed (GPF+, mCherry+, DeepRed-).

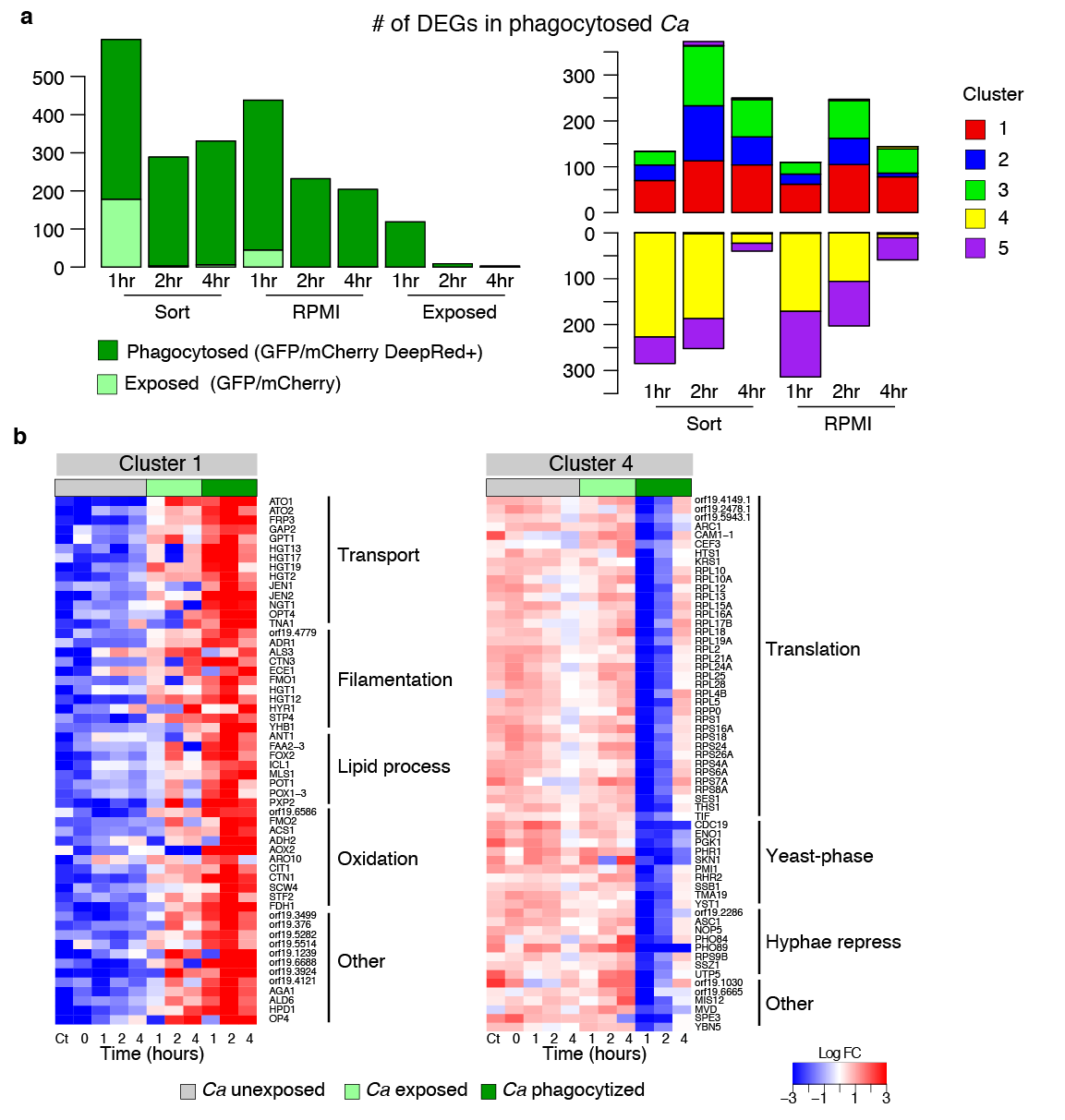

**Figure S2. Differential gene expression in phagocytosed *Candida albicans* subpopulation during macrophages infection. (a)** Number of genes differentially expressed (DEGs; FC >4, FDR < 0.001) in the live, phagocytosed subpopulation of *C. albicans* relative to exposed and unexposed (Sort, RPMI) subpopulations, and distribution of DEGs according the clusters of gene expression patterns. **(b)** Selection of most significantly DEGs from Cluster 1 (highly induced) and Cluster 4 (highly repressed) in *C. albicans* sorted populations (unexposed, exposed and phagocytosed) at 0, 1, 2 and 4 hours. Cluster 1 and 4 had the highest ratio of induction/repression upon phagocytosis. Synthesized functional biological categories were deducted from GO term enrichment and GO slim analyses (corrected-*p <* 0.05 hypergeometric distribution with Bonferroni correction).

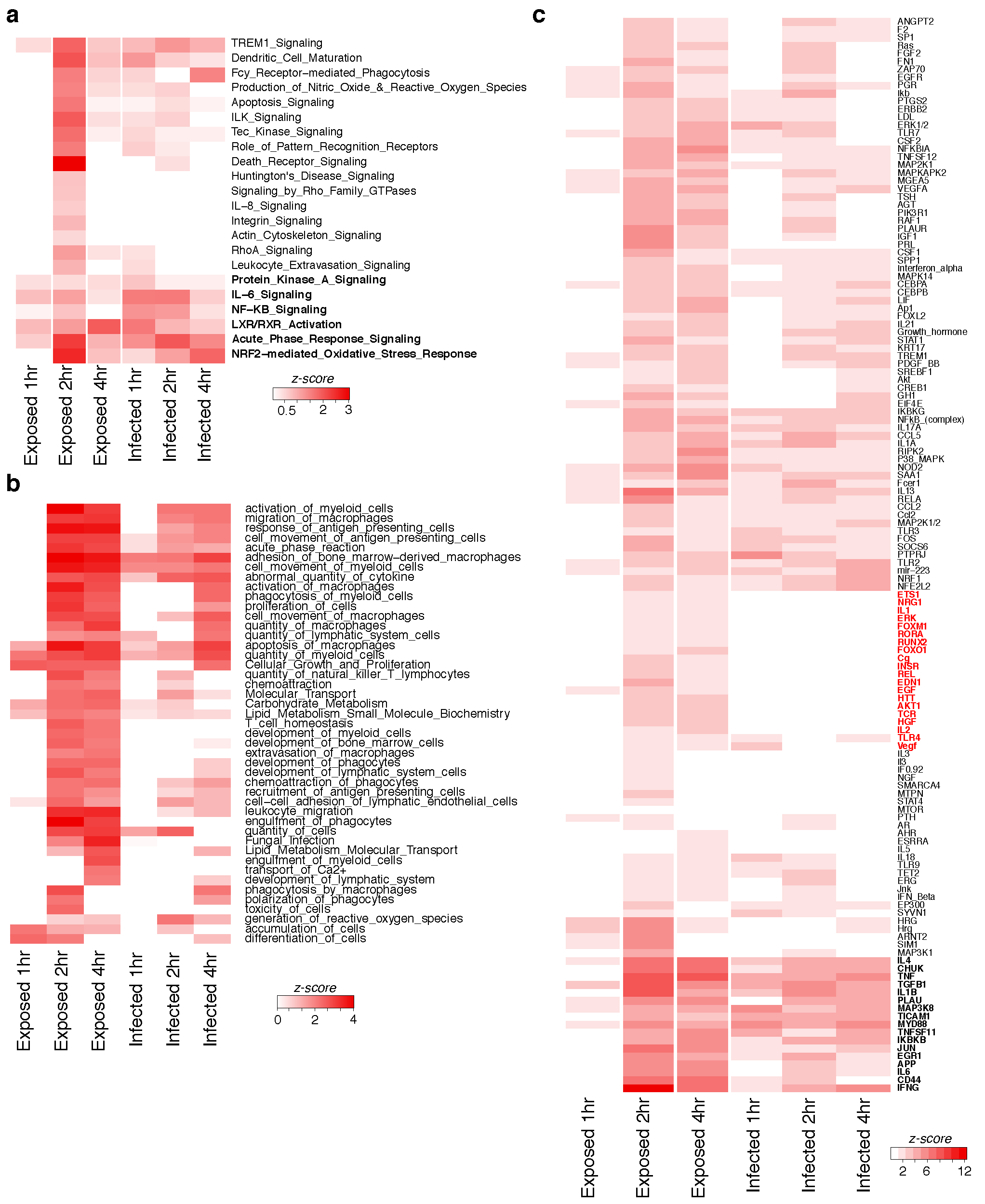

**Figure S3. Functional biological relationships of the transcriptional responses of macrophages to *Candida albicans* infection. (a)** Canonical pathways, **(b)** Biological functions, and **(c)** Upstream regulators induced in macrophages with phagocytosed *C. albicans* (Infected) and macrophages with non-phagocytosed fungus (Exposed) in comparison to macrophages unexposed to *C. albicans*. Categories in bold are significantly enriched in both infected and exposed macrophages

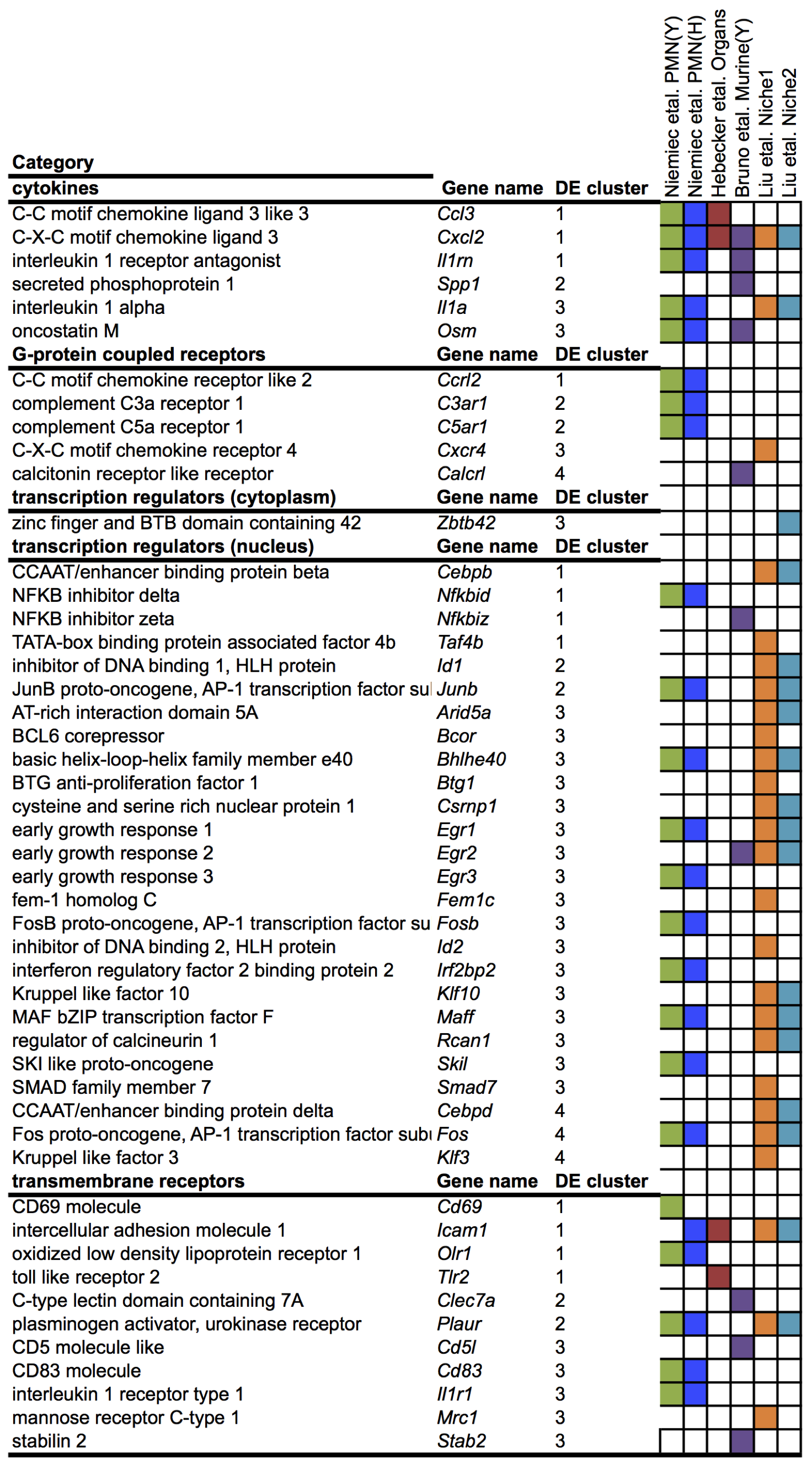

**Figure S4. Overall immune response genes to *Candida albicans* infection.** Comparison of immune response genes found significantly differentially expressed in this study and previously reported in other studies of the host transcriptional response to *C. albicans* interaction.

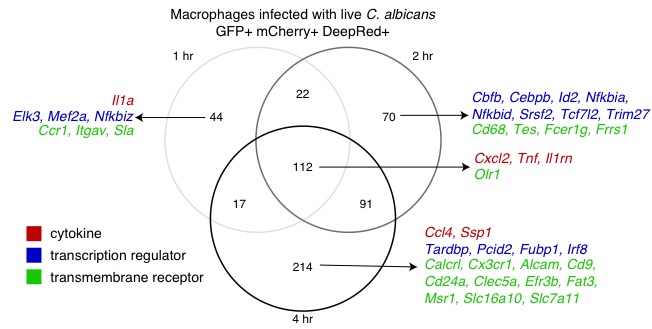

**Figure S5. Overlap of the transcriptional response in infected macrophages.** Different magnitude of induction for cytokines, transcriptional regulators, and transmembrane receptors differentially expressed in populations of macrophages infected with *Candida albicans* during 1, 2 and 4 hours post-infection.

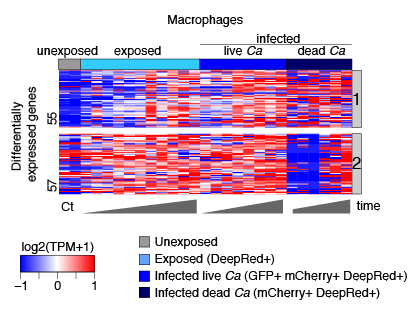

**Figure S6. Highly induced genes in subpopulation of macrophages infected with dead *Candida albicans*.** Heatmap depicts genes induced in subpopulation of macrophages infected with dead *Candida albicans* (clearance) relative to unexposed macrophages. Replicates for macrophages sorted subpopulations (unexposed, exposed and phagocytosed) at 0, 1, 2 and 4 hours post-infection are included. The gene expression is color coded from low expression (blue) to high expression (red) for each gene (row); two clusters of expression profile are shown (1 and 2, grey bars at right).

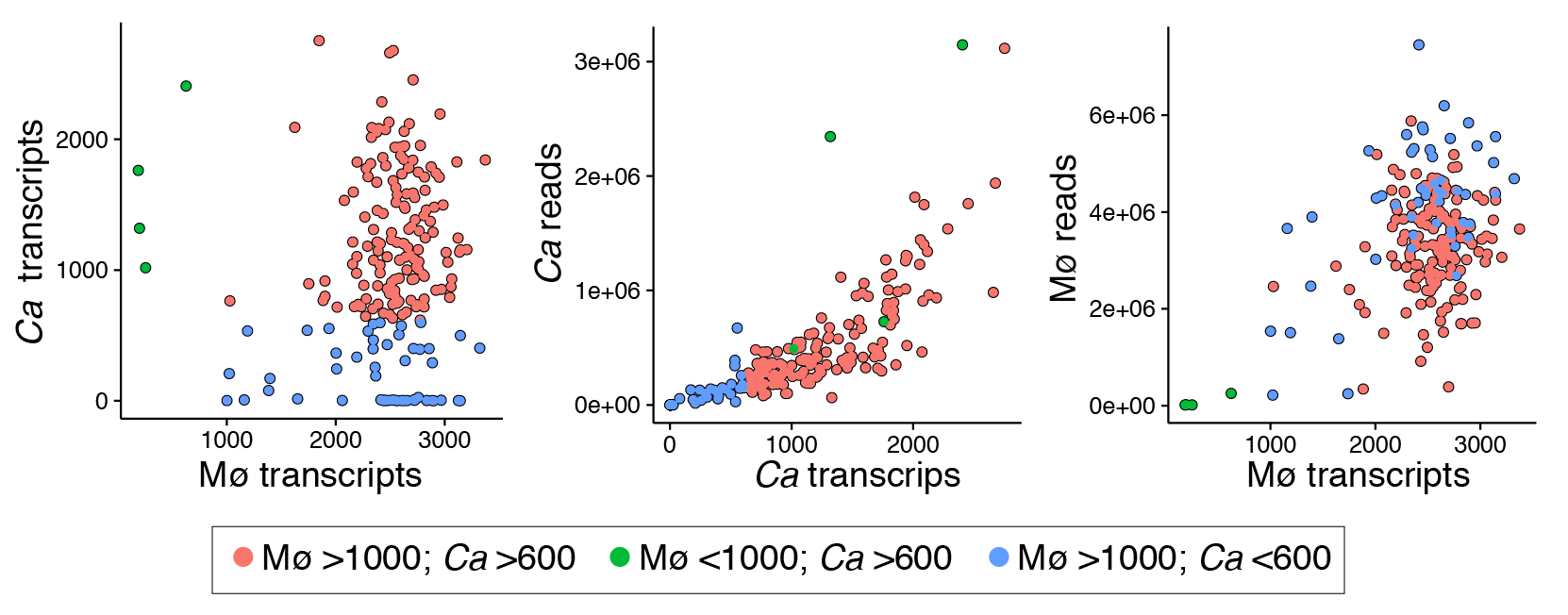

**Figure S7. Transcripts detected at single infected cell level in macrophages and phagocytized *C. albicans*.** Correlation plots of the number of detected transcripts (TPM > 1). Left: Comparison of the number of transcripts detected in macrophages and *C. albicans*. Middle and right: Comparison of transcripts detected (*x-axis*) and number of mapped reads (y-axis) for parallel infected macrophages with live *C. albicans* cells, respectively.

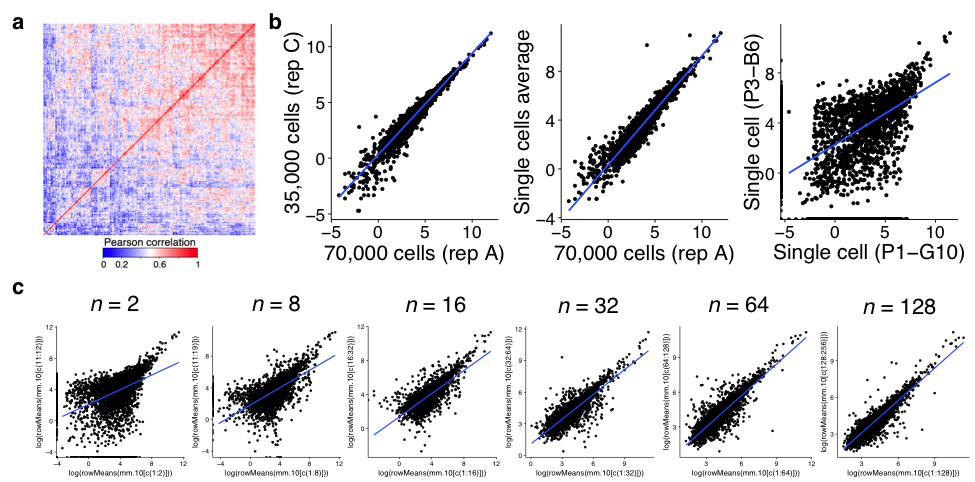

**Figure S8. Gene expression correlation between individual macrophages with phagocytized *C. albicans***. **(a)** Heat-map displays Pearson’s *r* values of the expression heterogeneity of the top 100 genes for 267 macrophages with phagocytized live *C. albicans* at 2 and 4 hrs. Red are highly correlated cells and blue are low correlated cells according the expression of the top 100 genes. **(b)** Plots of the gene expression correlation between two replicates of the sorted population of macrophages with phagocytized live *C. albicans* at 2 hr (left); between the single cells expression average and one replicate of the population (middle), and between two single cells (right). **(c)** Plots of the average gene expression correlation between increasing number of single-cells, *n* indicates the number of cells averaged.

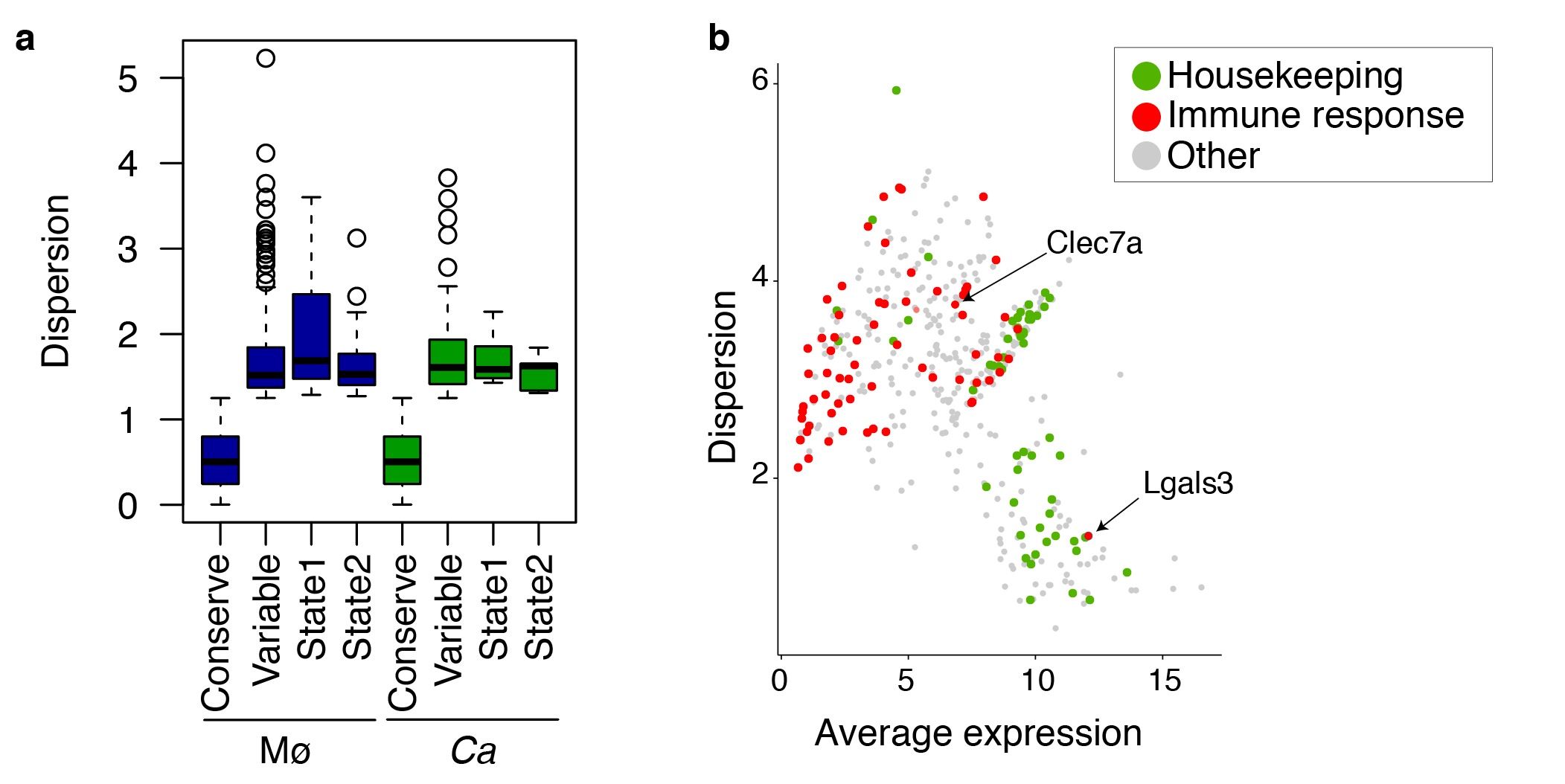

**Figure S9. Expression variability at the single-cell level. (a)** Boxplot of the dispersion of conserved and variable genes in macrophages infected and phagocytosed *C. albicans* for all genes, and for those in co-state1 and co-state2. **(b)** Relationship between average expression level in single macrophages with phagocytized *C. albicans* (*x axis*) and standard deviation (*y axis*) for ~3,000 genes. Red dots represent immune response genes, and green dots denote housekeeping genes. (**c**) Expression density distribution of five housekeeping genes across 267 macrophages with phagocytized live *C. albicans* cells at 2 and 4 hours. *y-axis*: cell density distribution; *x-axis*: single cell expression in log2(TPM +1) transformation.

**a**

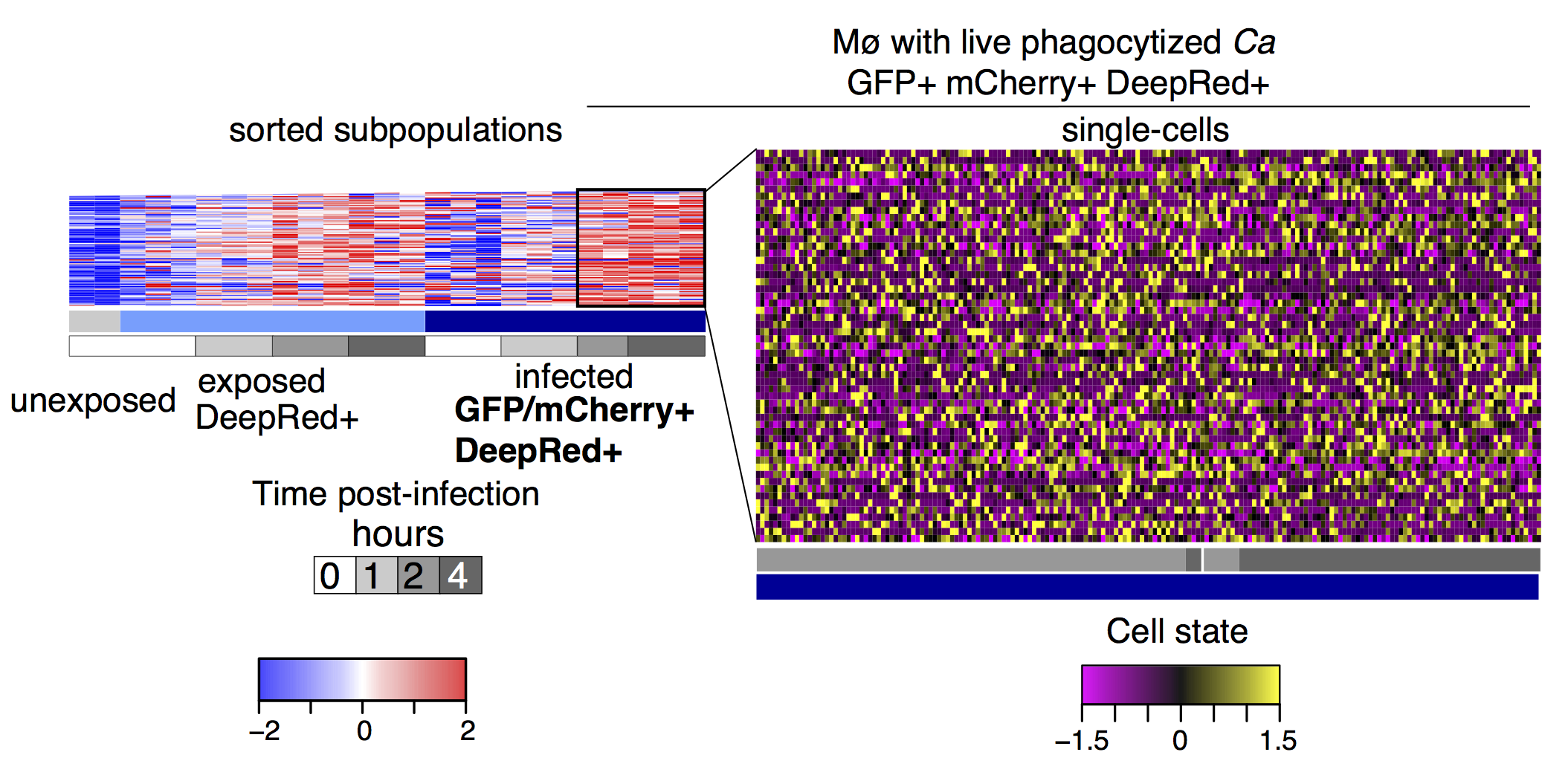

**b**

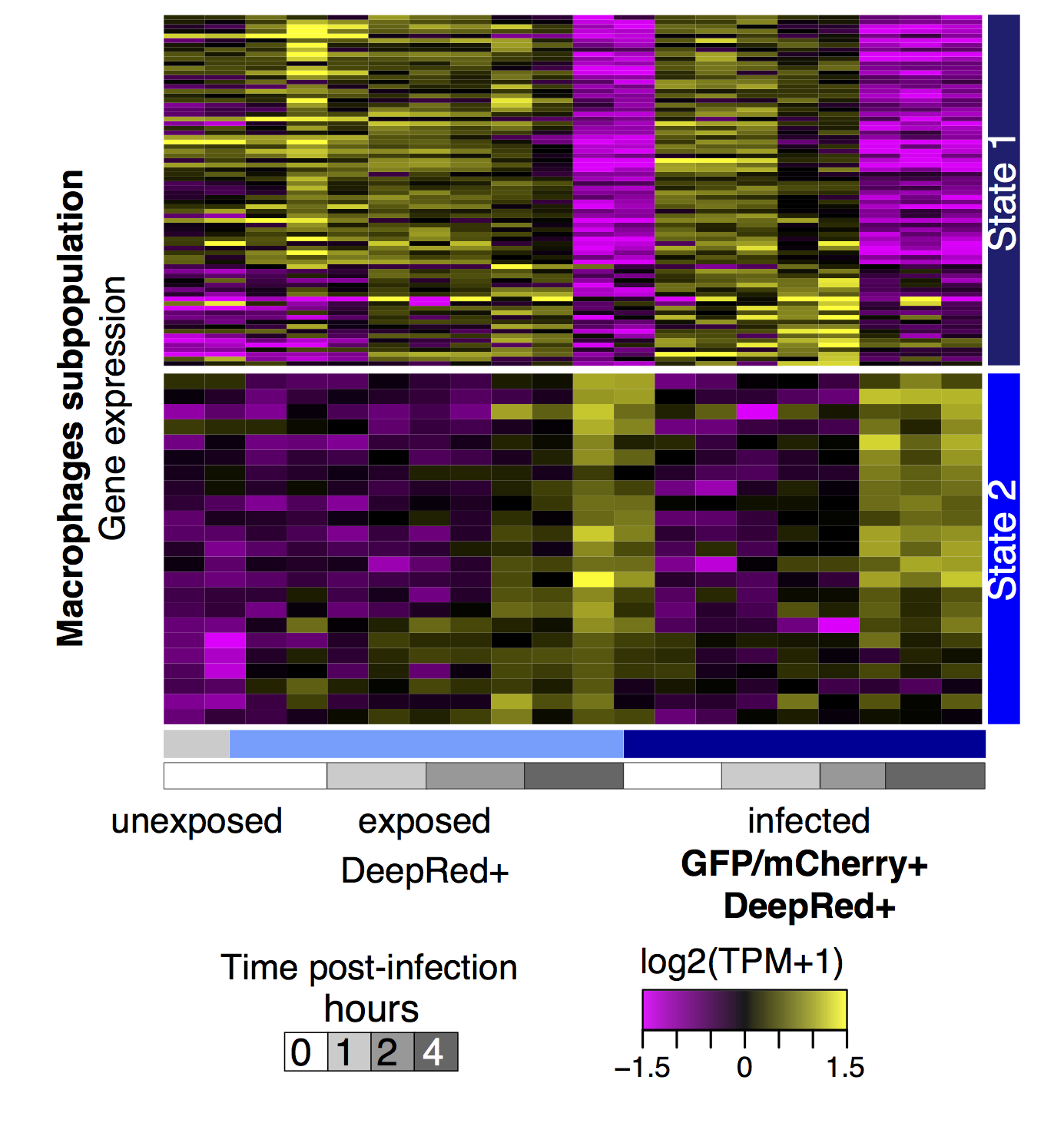

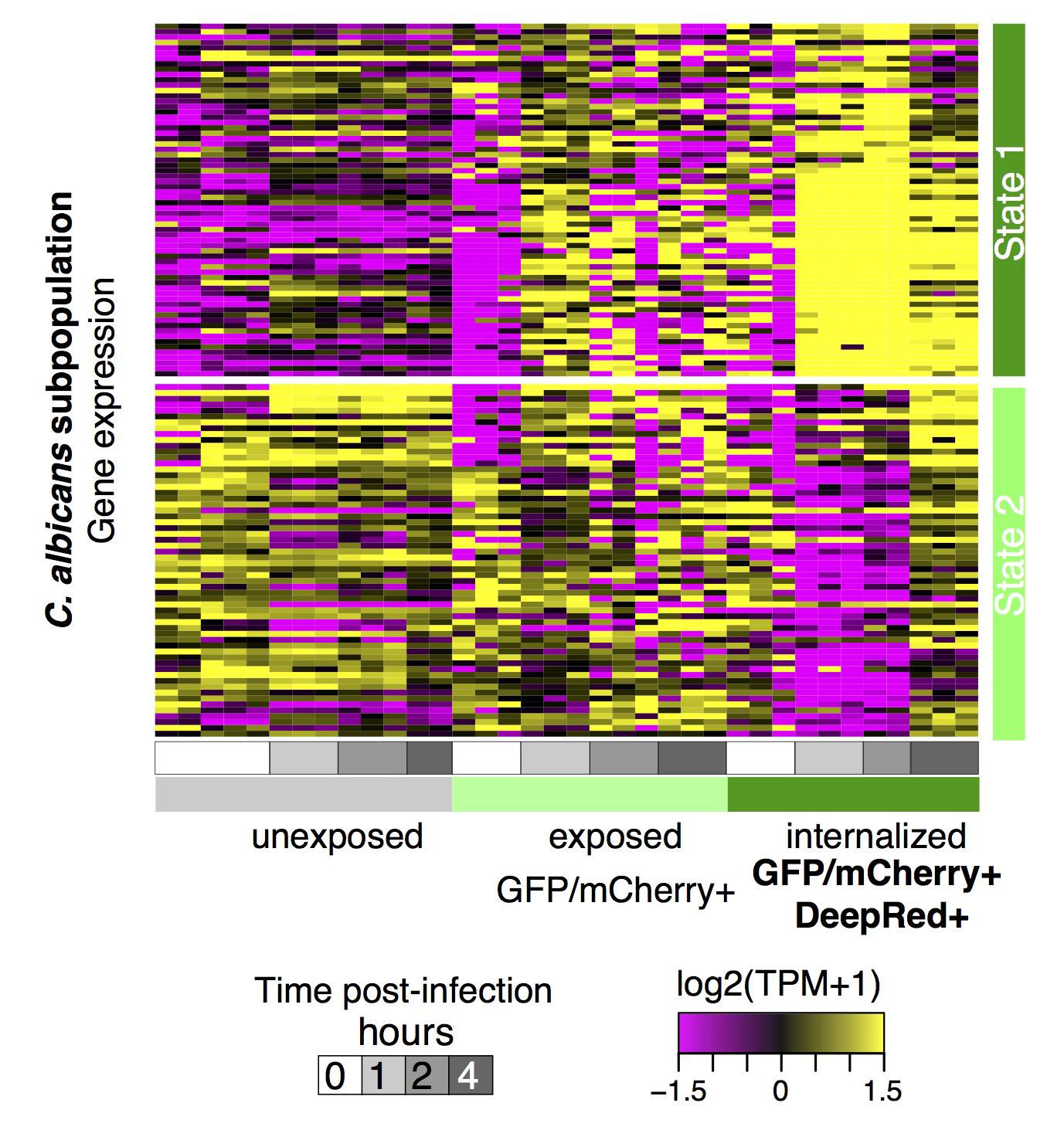

**Figure S10. Expression variability at the single-cell level of highly expressed genes in the population of live *C. albicans*-infected macrophages at 2 and 4 hours. (a)** Heatmap of DEGs in macrophages in sorted subpopulation samples (left) and expression of those genes in single infected macrophages (right). **(b)** Heatmap of the gene expression in the sorted subpopulations for the discriminative genes identified by single-cell analysis in co-state1 and co-state2 of single macrophages (left) infected with *C. albicans* (right).

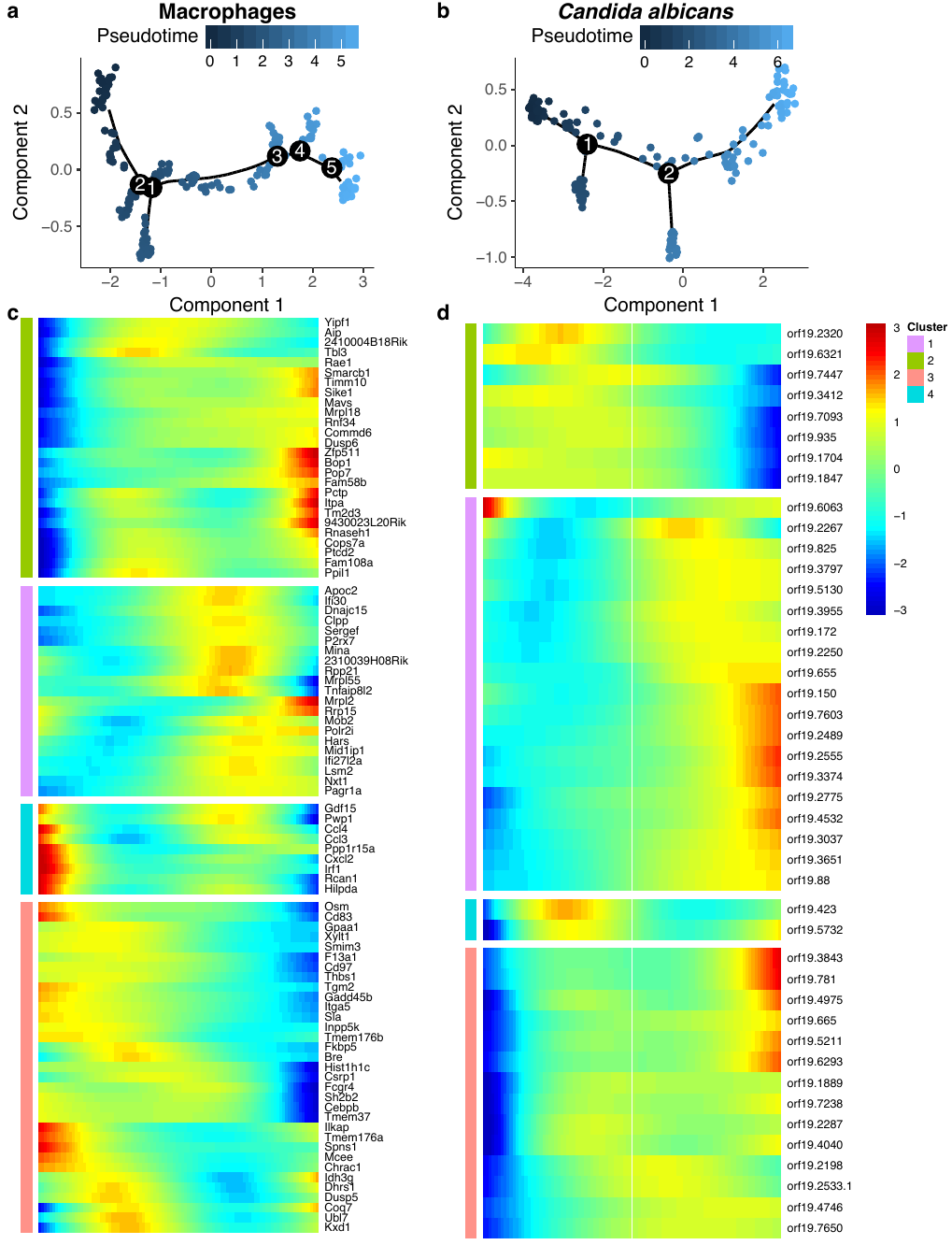

**Figure S11. Pseudo-time analysis in single infected macrophages and *C. albicans*.** tSNE plot for macrophages **(a)** and *C. albicans* **(b)** colored by the pseudo-time. Genes with most significant changes as a function of progress along the trajectory for **(c)** macrophages and **(d)** *C. albicans*.

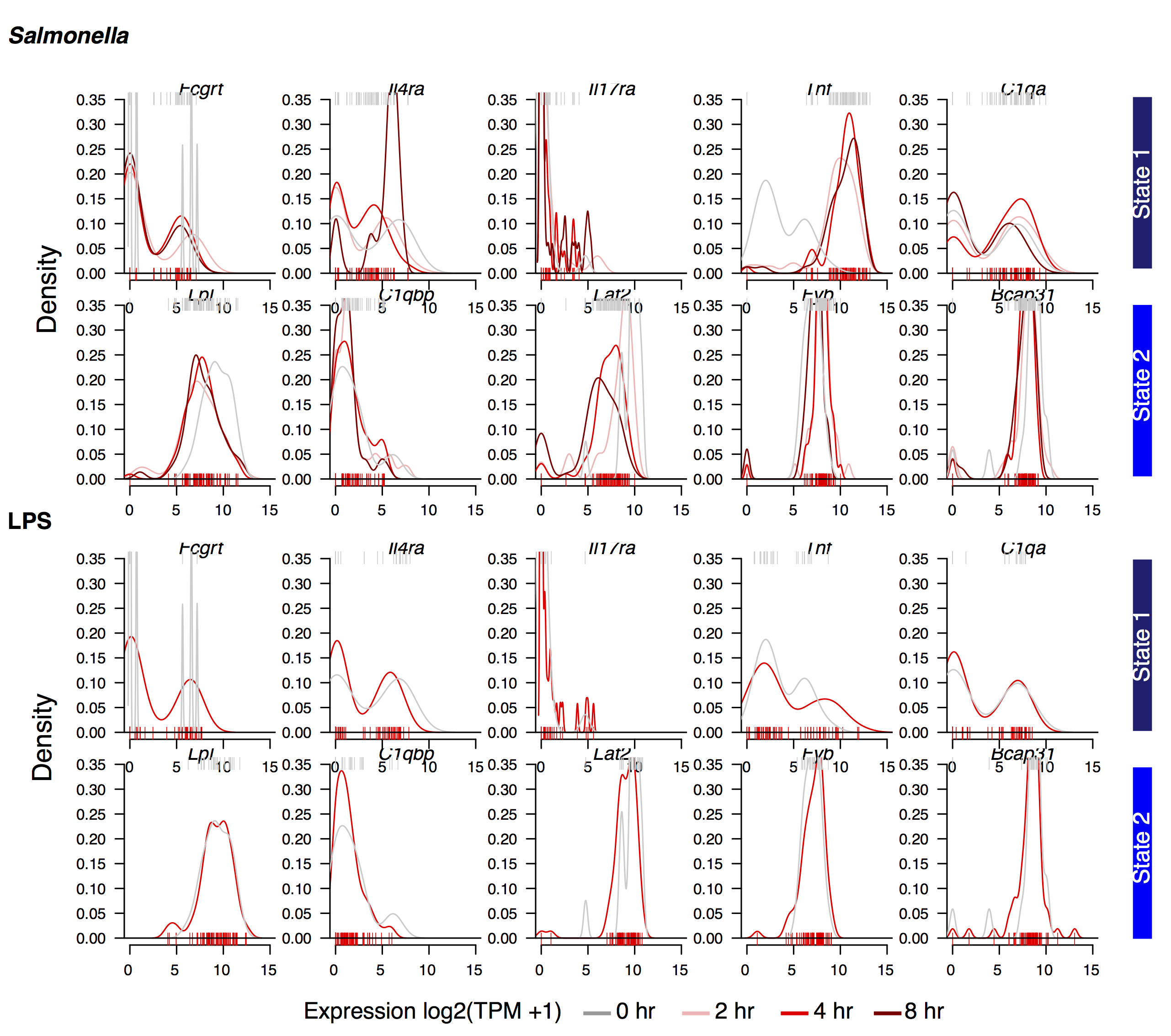

**Figure S12. Comparison with single-cell RNA-seq macrophages response to other stimuli.** Comparison of the expression density distributions for 5 top marker genes in co-state1 and co-state2 across macrophages-*C. albicans* single cells at 2 and 4 hours post infection (**Figure 5**) with the expression density of macrophages infected with *Salmonella* (top; *n* = 94) or macrophages exposed to LPS (bottom; *n* = 94) from Avraham, R., et al., 2015. Individual cells are plotted as bars underneath (red; 4 hours) or overhead (gray; 0 hours) for each distribution.

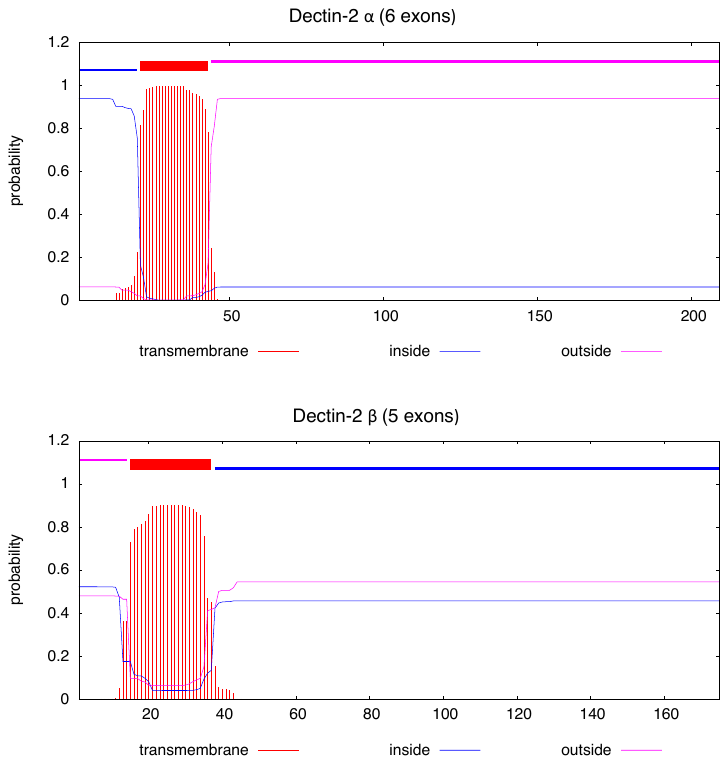

**Figure S13. Prediction of transmembrane helices in Dectin-2.** Prediction of transmembrane helices in the Dectin-2 α (Q9JKF4-1) and Dectin-2 β (Q9JKF4-2) amino acid sequences using TMHMM v2.0. and the hidden Markov model. The x -axis indicates the position in the amino acid sequence starting at the N-terminal end and the y -axis the probability of residing inside of, outside of, or in the membrane.

**Figure S14.** *Candida albicans* reporter strain displays reduced filamentation compared to SC5314. **(a)** SC5314, **(b)** CAI-F2 (reporter strain parent) and **(c)** CAI4-F2-*NEUT5L*-*NAT1*-*mCherry*-*GFP* strain (reporter strain) at 40x under identical growth conditions are phenotypically distinct. Fluorescent reporters were inserted into CAI-4-F2 **(b)** for our studies. Reduced filamentation made it possible to sort cells and monitor host interactions up to 4 hours at conditions that both mimic host environment *in vitro* and induce filamentation (RPMI medium, 10% fetal calf serum (not heat inactivated) at 37 °C and 5% C0_2_ for four hours). **(c)** The resulting strain, CAI-4-F2-*NEUT5L*-*NAT1*-*mCherry*-*GFP*, is phenotypically similar to the parent and was verified as being resistant to Nourseothricin and fluorescent for both GFP and mCherry.
